# Single-cell immune profiling of regional lymph nodes during early-stage breast cancer progression

**DOI:** 10.64898/2026.05.18.724563

**Authors:** Marit Otterlei Fjoertoft, Oystein Garred, Karin Teien Lande, Inger Riise Bergheim, Margit Hesla Riis, Ole Christian Lingjaerde, Hege Russnes, June Helen Myklebust, Kanutte Huse, Inga Hansine Rye

**Affiliations:** Dep.of Cancer Gentics, Institute for Cancer Research, Oslo University Hospital, Oslo, Norway; Dep.of Patology, Oslo University Hospital, Oslo, Norway; Dep.of Cancer Genetics, Institute for Cancer Research, Oslo University Hospital, Oslo, Norway; Dep.of Cancer Genetics, institute for Cancer Research, Oslo University Hospital, Oslo, Norway; Institute for informatics, University of Oslo, Oslo, Norway; Dep.of Cancer Immunology, institute for Cancer Research, Oslo University Hospital, Oslo, Norway; Dep.of Cancer Immunology, Institute for Cancer Research, Oslo University Hospital, Oslo, Norway; Institute for Cancer Research

## Abstract

**INTRODUCION:** Tumor cell infiltration in regional lymph nodes is a strong prognostic marker, guiding treatment decisions in breast cancer. While the immune cell composition in primary tumors has been more widely explored in later years, the immune cell composition of the sentinel node (SN) and axillary lymph nodes (ALN) remains understudied. A better understanding of how primary tumor and metastatic tumor cells alter the nodal immune microenvironment can shed light on metastasis and cancer progression to unveil new treatment strategies.

**MATERIALS AND METHODS:** From a prospective clinical cohort of 458 treatment-naive patients with primary operable breast cancer, we performed comprehensive immunophenotypic analysis using mass cytometry analysis of non-metastatic (SN-) and metastatic (SN+) and ALN (ALN+) lymph nodes.

**RESULTS:** As expected, patients with ALN+ cases had a shorter time to distant metastases than SN+ and SN- cases. We identified an exhausted T-cell phenotype and an increase in Germinal Center B (GC B) cells and plasma cells in ALN+ samples compared to SN- samples, both in the whole cohort as well as when investigating estrogen-receptor positive (ER^+^) patients only. There were no differences in immune cell composition across breast cancer (BC) subtypes within SN-samples. SN+ samples from triple negative BC (TNBC) showed a trend towards increased abundance of GC B and plasma cells, similar to more advanced ALN+, suggesting that smaller TN metastases may trigger an immune activation at an early stage of dissemination. Further analysis of SN- samples from ER^+^ patients revealed a subset of patients where the immune response had a more exhausted T-cell phenotype. This group was enriched for lymph nodes that were deemed negative by ordinary pathology examination (microscopy) but had detectable tumor cells by CyTOF analysis.

**CONCLUSION:** The immune profiles of SN and ALN samples from breast cancer patients are highly diverse, showing limited associations to BC subtype, clinical parameters or patient outcome. Metastatic tumor cells play a significant role in driving T-cell exhaustion and immunosuppression. Notably, in approximately 50% of the ER^+^ samples, T-cell exhaustion was detectable. This coincides with the presence of tumor cells identified by CyTOF, which were likely missed by conventional pathological examination. These findings suggest that small tumor deposits alter the immune composition, and the immune profile reveals the presence of tumor cells.

## Introduction

Breast cancer, the most common cancer among women worldwide (1), is a heterogeneous disease, with prognosis and treatment strategies dependent on tumor subtype, histological grade, lymph node status, and disease stage. Histological subtyping is based on the expression of estrogen receptor (ER), progesterone receptor (PgR), and human epidermal growth factor receptor 2 (Her2). ER^+^ tumors comprise approximately 70% of cases, followed by Her2+ (14%) and triple-negative breast cancer (TNBC, 11%) (2). Surgery, chemotherapy, and radiation, among others, are treatment options provided to all breast cancer patients. In addition, patients with ER- and/or PgR+ tumors receive endocrine therapy, while those with Her2+ disease are treated with Her2-targeted therapies in addition to standard chemotherapy (3). TNBC, characterized by the highest levels of immune infiltration (4), has shown a promising response to the PD-L1 immune checkpoint inhibitor pembrolizumab when combined with chemotherapy in PD-L1+ cases (5,6).

Lymph node status is an independent prognostic factor, with tumor size, grade, age, and subtype predicting nodal involvement (7). Removal and examination of the sentinel node is the standard of care for staging in patients with early breast cancer and in selected cases of locally advanced breast cancer. The guidelines on the surgical procedure in the axilla, the possible radiation of the axilla, and further systemic therapy are continuously changing according to results from the most recent publications (8). They are based upon results from histological examination of lymph nodes (pN) in addition to clinical nodal status (cN), either detected by radiological examination or palpation (9–13). In cases with more advanced disease in the axilla, a complete axillary dissection is performed without prior SN procedure (11). Larger metastases (>2 mm), higher number of affected nodes and perinodal growth are linked with poorer outcomes (14,15), and these adverse features are most commonly observed in patients with ER^+^ and Her2^+^ tumors (15).

The presence of lymph node metastases is a well-established indicator of increased risk of distant metastasis. However, whether lymph nodes facilitate further dissemination to distant sites such as lungs, bone marrow and brain remains debatable. While some studies, such as Ullah *et al.* (16), suggest that axillary lymph node metastases may not directly contribute to seeding of distant metastasis, contrasting evidence exists. Naxerova *et al.* (17) and Gundem *et al.* (18), using colon and prostate cancer, reported that a significant portion of distant metastases shared the same subclonal origin as the lymph node metastasis. Additionally, studies in mice have provided interesting insights. Reticker-Flynn and colleagues (19) demonstrated that lymph node colonization by tumor cells can induce tumor-immune tolerance by the systemic increase of regulatory T cells (Tregs), facilitating distant metastasis. Similarly, Brown *et al.* (20) and Pereira *et al.* (21) showed the ability of a TNBC mouse cell line to egress from the lymph node and colonize the lungs. These findings collectively highlight the multifaceted role of lymph nodes in tumor progression. Lymph nodes serve not only as transit sites for disseminating cancer cells but also potentially contribute to the establishment of a favorable microenvironment for distant metastasis.

Recent single-cell RNA sequencing studies have compared the immune cell landscape in primary tumors and metastatic lymph nodes (22,23). However, how early-stage breast cancer progression alters the nodal immune cell landscape remains unclear. We have previously shown in a smaller study that metastatic nodes have a shift in the immune cell landscape towards an increase in memory and exhausted T cells and Tregs (24,25). The change in the abundance of T cell subsets was correlated with tumor burden (24,25). These studies were, however, not designed to address potential BC subtype differences. In this study we conducted a comprehensive immune cell profiling of non-metastatic and metastatic SN and ALN from 458 treatment-naive breast cancer patients, encompassing ER^+^, Her2^+^ and TNBC subtypes.

## Materials and methods

### Sample collection

SN and ALN specimens were collected from 466 patients with operable breast cancer (T1-T2) and without distant metastasis from the OSLO2 breast cancer observational trial between the years 2010 and 2016. Following surgery, patients received chemotherapy, endocrine therapy and/or radiation therapy according to the national guidelines (3). The samples were collected after written patient consent in accordance with the Declaration of Helsinki and approved by the regional committee for research ethics (200606181-1, 538-07278a, 2009/4935, 2016/433). Clinical data were collected from the Cancer Registry of Norway, with updates in distant metastasis and death status obtained as of September 2024. In Norway, the standard procedure for SN detection in breast cancer involves subcutaneous periareolar injection of the radioactive tracer technetium-99m and methylene blue dye. The first lymph nodes to absorb either the dye or radioactivity, usually one or two, are identified as the sentinel nodes. At the time of this study, a complete axillary metastasis was performed in cases with known metastasis preoperatively or confirmed in SN, and the threshold for doing an axillary dissection was lower than what the current national guideline suggests (3). Both the SN and ALN were removed during surgical excision of the primary tumor.

The lymph nodes were further examined by standard pathology examination where one half of each node were processed as formalin-fixated, paraffine-embedded (FFPE) tissue, sectioned, stained by haematology and eosin (HE) and by microscopy classified into three categories based on metastatic tumor burden and location: non-metastatic SN (SN-, *n* = 362), metastatic SN (SN+, *n* = 90), and metastatic ALN (ALN+, *n* = 13). Sixteen of the SN- samples were sampled from patients with non-invasive primary tumors, ductal carcinoma *in situ* (DCIS). The SN+ samples were further stratified by metastatic burden according to guidelines (7): SN+ with isolated tumor cells (SN+ITC, <0.2 mm or <100 cells, *n* = 7), SN+ with micrometastasis (SN+micromet, 0.2-2 mm, *n* = 18) and SN+ with macrometastasis (SN+macromet, >2 mm, *n* = 65). Primary tumors were evaluated for standard parameters including histological grade and protein markers (ER, PgR, Her2 and Ki-67) following ASCO/CAP recommendations (26) (Table 1). Eight samples were excluded based on exclusion criteria such as live cell count <65%, multiple samples per patient and samples lacking pathological information, resulting in a final cohort of 458 lymph nodes (Supplementary Fig. 1). The ALN+ samples were included in prior publication (25).

**Table 1:**
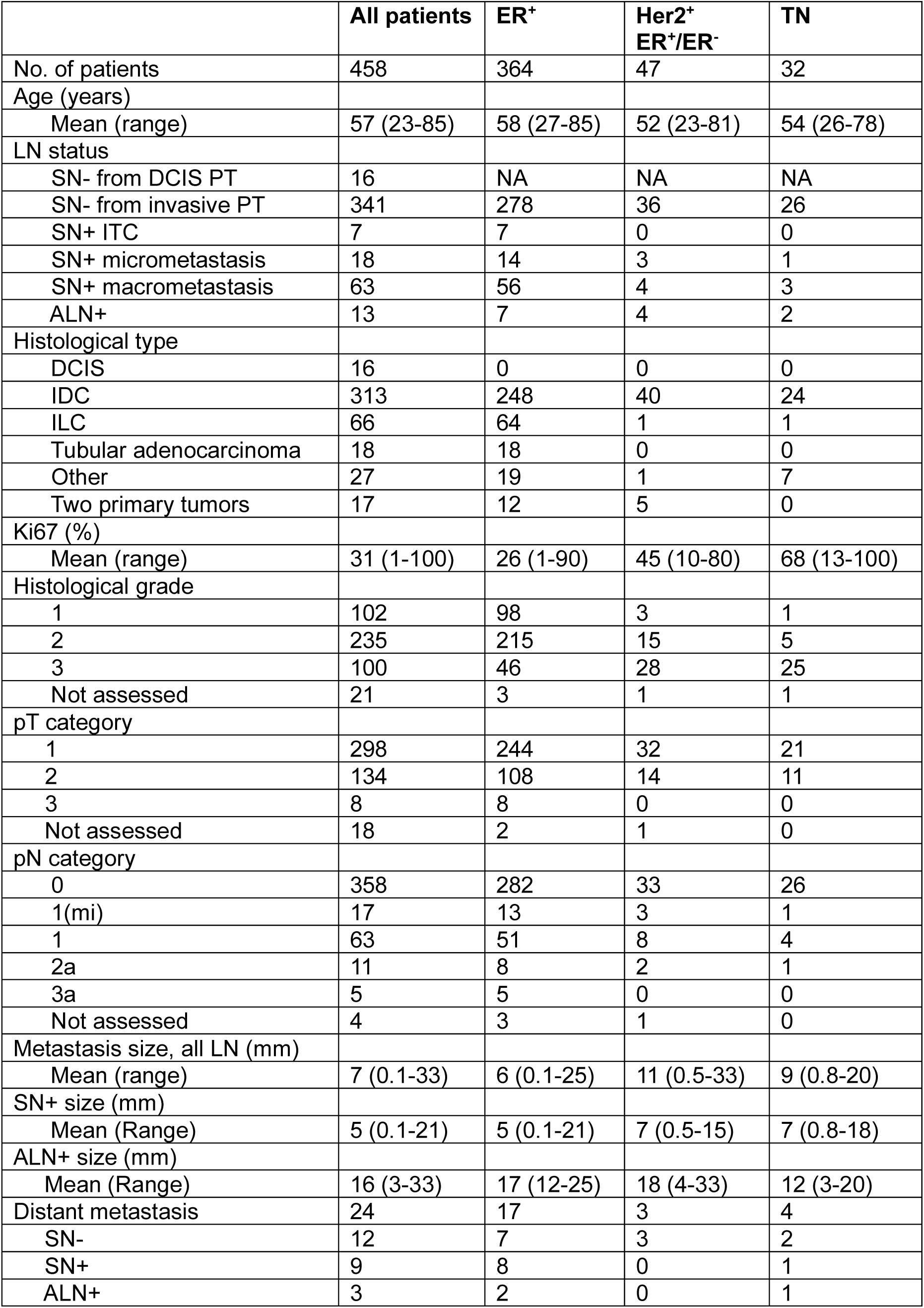
Clinical and histopathological parameters of the cohort after sample filtering (*n* = 458). LN = lymph node, ITC = isolated tumor cells, PT = primary tumor, DCIS = ductal carcinoma *in situ*, IDC = invasive ductal carcinoma, ILC = invasive lobular carcinoma.

|  | All patients | ER<sup>+</sup> | Her2<sup>+</sup><br>ER<sup>+</sup>/ER<sup>-</sup> | TN |
| --- | --- | --- | --- | --- |
| No. of patients | 458 | 364 | 47 | 32 |
| Age (years) |  |  |  |  |
| Mean (range) | 57 (23-85) | 58 (27-85) | 52 (23-81) | 54 (26-78) |
| LN status |  |  |  |  |
| SN- from DCIS PT | 16 | NA | NA | NA |
| SN- from invasive PT | 341 | 278 | 36 | 26 |
| SN+ ITC | 7 | 7 | 0 | 0 |
| SN+ micrometastasis | 18 | 14 | 3 | 1 |
| SN+ macrometastasis | 63 | 56 | 4 | 3 |
| ALN+ | 13 | 7 | 4 | 2 |
| Histological type |  |  |  |  |
| DCIS | 16 | 0 | 0 | 0 |
| IDC | 313 | 248 | 40 | 24 |
| ILC | 66 | 64 | 1 | 1 |
| Tubular adenocarcinoma | 18 | 18 | 0 | 0 |
| Other | 27 | 19 | 1 | 7 |
| Two primary tumors | 17 | 12 | 5 | 0 |
| Ki67 (%) |  |  |  |  |
| Mean (range) | 31 (1-100) | 26 (1-90) | 45 (10-80) | 68 (13-100) |
| Histological grade |  |  |  |  |
| 1 | 102 | 98 | 3 | 1 |
| 2 | 235 | 215 | 15 | 5 |
| 3 | 100 | 46 | 28 | 25 |
| Not assessed | 21 | 3 | 1 | 1 |
| pT category |  |  |  |  |
| 1 | 298 | 244 | 32 | 21 |
| 2 | 134 | 108 | 14 | 11 |
| 3 | 8 | 8 | 0 | 0 |
| Not assessed | 18 | 2 | 1 | 0 |
| pN category |  |  |  |  |
| 0 | 358 | 282 | 33 | 26 |
| 1(mi) | 17 | 13 | 3 | 1 |
| 1 | 63 | 51 | 8 | 4 |
| 2a | 11 | 8 | 2 | 1 |
| 3a | 5 | 5 | 0 | 0 |
| Not assessed | 4 | 3 | 1 | 0 |
| Metastasis size, all LN (mm) |  |  |  |  |
| Mean (range) | 7 (0.1-33) | 6 (0.1-25) | 11 (0.5-33) | 9 (0.8-20) |
| SN+ size (mm) |  |  |  |  |
| Mean (Range) | 5 (0.1-21) | 5 (0.1-21) | 7 (0.5-15) | 7 (0.8-18) |
| ALN+ size (mm) |  |  |  |  |
| Mean (Range) | 16 (3-33) | 17 (12-25) | 18 (4-33) | 12 (3-20) |
| Distant metastasis | 24 | 17 | 3 | 4 |
| SN- | 12 | 7 | 3 | 2 |
| SN+ | 9 | 8 | 0 | 1 |
| ALN+ | 3 | 2 | 0 | 1 |

### Dissociation protocol

Half of the lymph node was processed and dissociated into single-cell suspension and cryopreserved as previously described (24,25), including adding collagenase, hyaluronidase and BSA during the processing of the ALN+ samples (24).

### Mass cytometry and analysis of mass cytometry data

Mass cytometry analysis of the cryopreserved single-cell samples was performed as previously described (Supplementary Table 1) (25). Shortly summarized, the cryopreserved cells were thawed and stained using surface and intracellular antibody cocktails frozen in aliquots and kept at −80 °C until use to minimize antibody variability. A control sample, consisting of peripheral blood mononuclear cells (PBMC) spiked with equal amounts of the two breast cancer cell lines BT-474 (ATCC®HTB-20™) and MDA-MB-231 (ATCC®HTB-26™), was included in each experimental run. The data were normalized with EQ beads using the CyTOF® Software 7.0 (Standard BioTools) version 7.0.8493, and the cloud-based analysis platform Cytobank was used for further analyses and expert gating. For semi-automatic cell type identification, we utilized a sequential FlowSOM approach, with a cleanup FlowSOM (FlowSOM 1) and a cell identification FlowSOM (FlowSOM 2), designed and described in previous publication (Supplementary Table 1) (25). Due to size limitations in Cytobank, FlowSOM 1 and FlowSOM 2 were each performed as 45 parallels runs, using the same SOM and settings across each run to ensure reproducibility. Due to a high number of tumor-negative samples in this cohort, tumor cells were not included in the FlowSOM 2 analysis. Cells identified as tumor, apoptotic and unclassified cells were manually evaluated and excluded from further analysis, resulting in 33 immune metaclusters (MCs). MCs were manually annotated into cell types based on marker expression, and MCs of very similar cell types were merged, giving a total of 27 immune cell populations (Supplementary Fig. 2). Tumor cell counts were manually curated and exported from FlowSOM 1 (Supplementary Fig. 1).

### Statistics

The statistical programming language R was used for boxplots, linear regressions analyses, hierarchical consensus clustering and statistical testing. All box plots indicate a box of the upper and lower quartile with a median value line, and whiskers indicating maximum and minimum data values falling within 1.5 times the interquartile range. The non-parametric Mann-Whitney *U* test was performed to calculate the statistical significance between two sample groups. All p-values were adjusted for multiple testing using Bonferroni correction using 27 groups, unless otherwise noticed. Cytobank was used for opt-SNE visualization and FlowSOM clustering (27).

Hierarchical consensus clustering was performed using cell type fractions, where columns are normalized by total CD45+ cells per sample, and rows are normalized across all samples. Both are clustered by Pearson correlation distance matrix and average linkage.

Survival curves were generated using the Kaplan-Meier method to assess distant disease-free survival (DDFS). DDFS was defined as the time in months from diagnosis to development of distant metastasis. Patients who had not experienced metastasis by the time of the last follow-up were censored. Differences in survival between patient groups were compared using the log-rank test. A p-value of <0.05 was considered statistically significant.

## Results

### Immune profiles of local lymph nodes from breast cancer patients

We performed large-scale mass cytometry profiling of lymph nodes from 458 breast cancer patients. Of these, 362 had primary tumors classified as ER^+^, 47 as Her2^+^, 32 as triple negative (TNBC) and 16 as ductal carcinoma *in situ* (DCIS). All patients had primary operable breast cancer without distant metastasis at the time of diagnosis, but varying metastatic status of regional lymph nodes as reviewed by pathological examination: non-metastatic SN (SN-, *n* = 357), metastatic SN (SN+, *n* = 88) and metastatic ALN (ALN+, *n* = 13) (Fig. 1A, Table 1).

**Figure 1:**
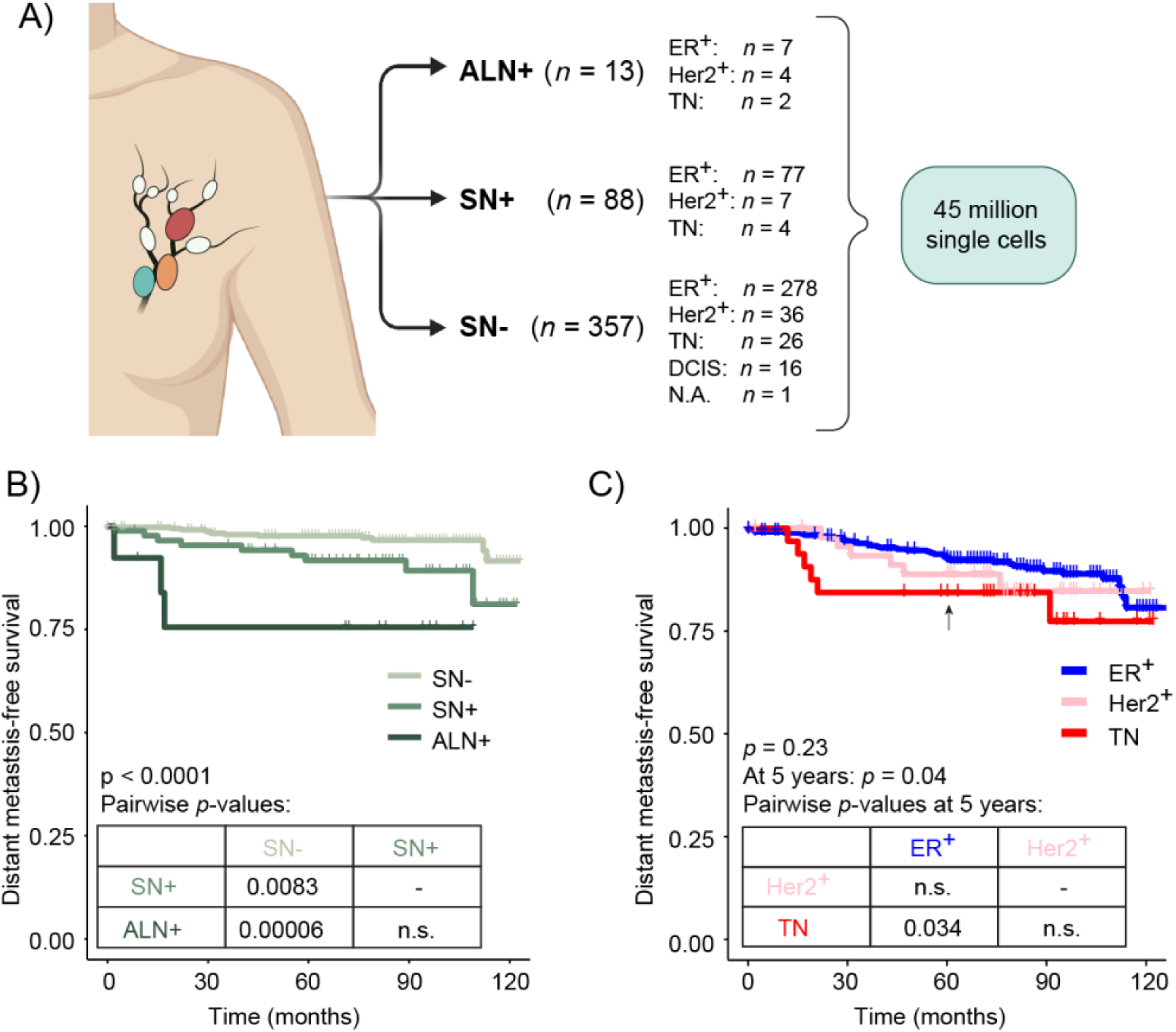
Overview of the cohort of lymph nodes from 458 breast cancer patients. **A)** Overview of the breast cancer cohort, the number of patients within each nodal category and their primary tumors’ main characteristics. **B)** Kaplan-Meier curves for distant disease-free survival (DDFS), defined as time to distant metastasis. Patients were stratified according to lymph node metastatic status, and **C)** according to breast cancer subtypes. 5-years follow-up is indicated by an arrow. Pairwise log-rank tests with Benjamini-Hochberg post-hoc correction were performed on the survival analyses. SN-: sentinel lymph node without metastasis, SN+: metastatic sentinel lymph node, ALN+: metastatic axillary lymph node.

A total of 24 (5.2%) patients developed distant metastasis during the time of follow-up (Table 1). Patients with ALN+ had the highest risk of developing distant metastases, showing significantly shorter DDFS compared to patients with negative nodal status (*p* = 0.000006, Fig. 1B). The risk of distant metastasis for patients with ALN+ was most significant within the first two years following diagnosis. Patients with SN+ also exhibited a higher risk for progression of the disease compared to SN- patients (*p* = 0.0083, Fig. 1B). The 10-year risk for distant metastases was not significantly different across BC subtypes, however, the TN subtype showed worse 5-year DDFS compared to the ER^+^ subtype (*p* = 0.034) (Fig. 1C), as also reported by others (15).

By using a comprehensive 46-parameter antibody panel, we analyzed a total of 45 million single cells and by FlowSOM analysis we identified 27 subpopulations of B cells, T cells, natural killer (NK) cells and myeloid cells (Fig. 2A, Supplementary Fig. 2, Supplementary Table 1). Visualization of all cell types in an opt-SNE run on all lymph nodes revealed a separation of the main cell types into distinct islands (Fig. 2A, Supplementary Fig. 3). The composition of the main cell types varied across SN-, SN+ and ALN+ to a similar degree for each BC subtype (Fig. 2B). The median frequency of the main cell types was similar across BC subtypes when viewed across all lymph nodes (Fig. 2C).

**Figure 2:**
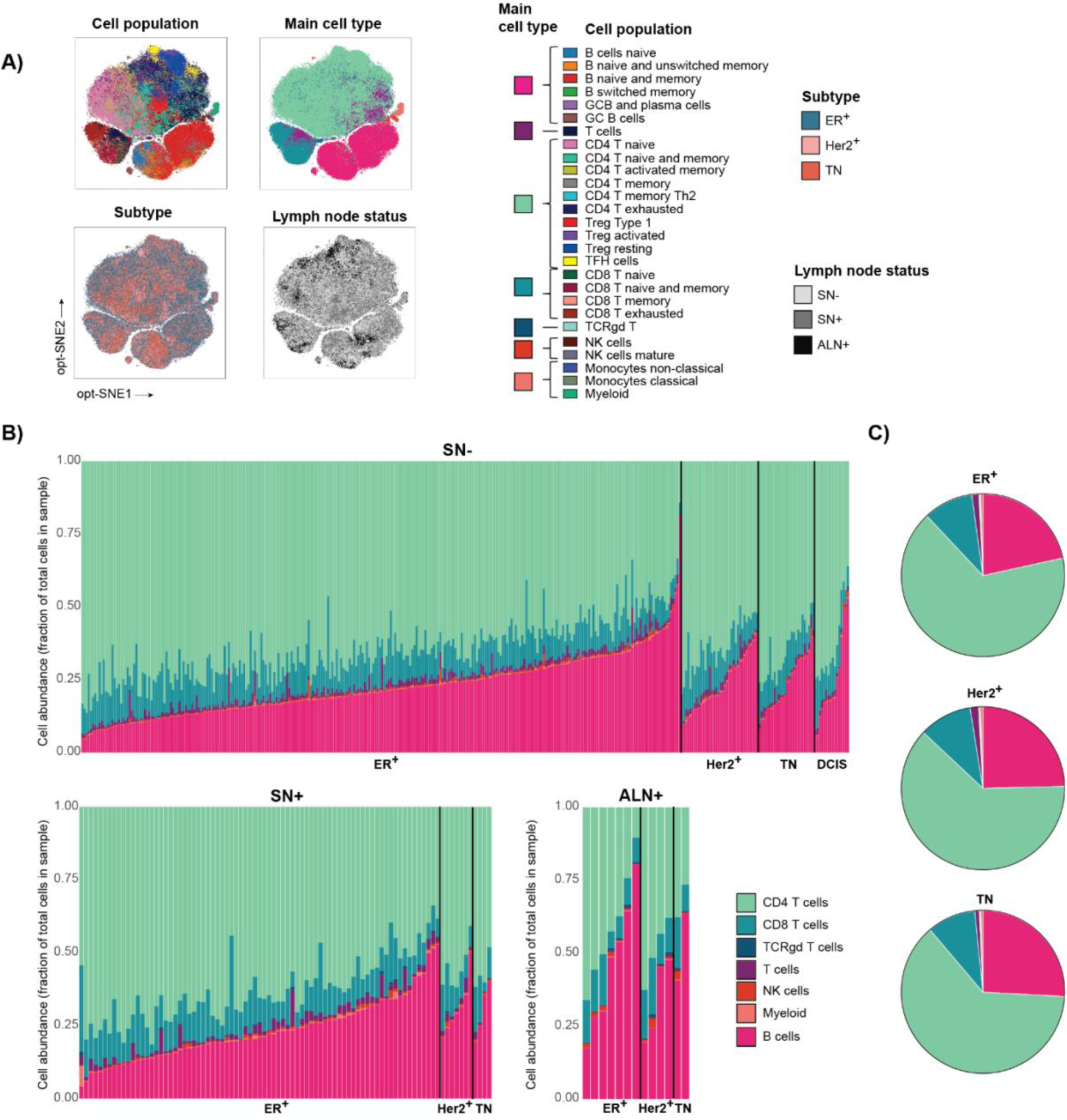
Identification of cell types in local lymph nodes. Single-cell mass cytometry (CyTOF) was used to analyze the lymph node samples. **B)** Opt-SNE showing the distribution of the 27 immune cell populations, main cell types, lymph node status and BC subtypes (ER^+^, Her2^+^, TN), *n* = 458. **C)** Stacked bar charts for SN-, SN+ and ALN+ of the distribution of the main cell types across the different lymph node sample types, and herein sorted by the different subtypes, and overview of median immune cell abundance of main cell types within the different breast cancer subtypes.

### Increased abundance of B cells and exhausted T cells in ALN+ samples

A more distinct difference in immune-phenotype between ALN+ and SN+/SN-samples was identified by a more fine-grained plot of all 27 immune cell populations. Naive CD4 T cells were the largest population in SN- and SN+ samples, whereas naive and memory B cells were the largest population in ALN+ samples (Fig. 3A).

**Figure 3:**
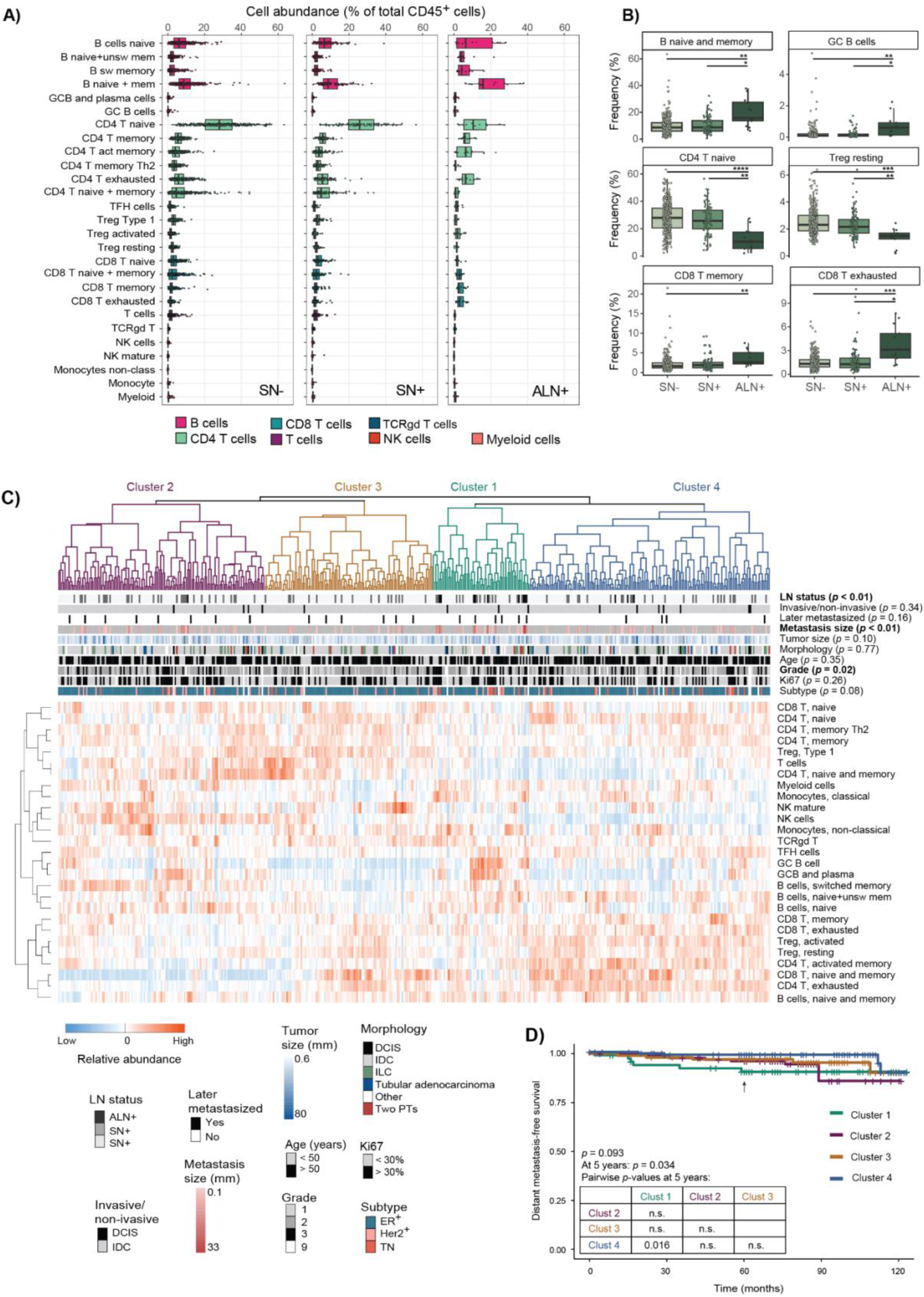
Immune cell profiles of sentinel and axillary lymph nodes show increased memory and exhaustion CD8 T cells and B cell abundance in ALN+ samples. **A)** Abundance of immune cells (27 cell populations) colored by main cell type in the three lymph node statuses. **B)** The frequencies of six selected immune cell populations across SN-, SN+, and ALN+ samples (Mann Whitney, Bonferroni corrected for 27 tests: \**p* < 0.05, \*\**p* < 0.01, \*\*\**p* < 0.001, \*\*\*\**p* < 0.0001). Remaining immune cell populations are shown in Supplementary Figure 4 with *p*-values in Supplementary Table 2. **C)** Hierarchical consensus clustering using cell type fractions for all lymph node samples (*p*-values from Fisher’s exact test). Columns are normalized by total CD45+ cells per sample, and rows are normalized across all samples. Both are clustered by Pearson correlation distance matrix and average linkage. **D)** Kaplan-Meier curves showing distant disease-free survival (DDFS) with patients stratified according to cluster identity. 5-year follow-up indicated by an arrow. Pairwise log-rank tests with Benjamini-Hochberg post-hoc correction were performed on the survival curves.

The ALN+ samples had significantly higher abundance of memory and exhausted CD8 T cells and lower abundance of naive and memory CD4 T cells and Treg Type 1 compared to SN+/SN- samples (Fig. 3B, Supplementary Fig. 4), which is in line with previous findings (24,25). Additionally, the frequencies of naive, unswitched and switched memory B cells, GC B cells, monocytes and resting Tregs were significantly increased in ALN+ samples compared to SN+/SN- samples (Fig. 3B, Supplementary Fig. 4). In concordance with previous work (24,25), we found no significant differences in immune cell phenotypes between SN- and SN+ samples but noted large variability in cell population frequencies within the SN- and SN+ samples. (Fig. 3B, Supplementary Fig. 4).

To further investigate differences in immune cell composition during tumor progression, hierarchical consensus clustering of all 458 lymph nodes based on the cell type frequency was performed. Interestingly, 9 of the 13 ALN+ samples were in one cluster (Cluster 1 (green)), while the SN- and SN+ samples were distributed across all clusters (Fig. 3C). The ALN+ enriched cluster (Cluster 1) was characterized by high abundance of GC B cells and plasma cells and a lower abundance of naive and memory CD4 T cells. The cluster was also significantly correlated with a larger size of the metastasis and showed a trend towards more Her2^+^ and TNBC subtypes. Additionally, this cluster had a significantly lower DDFS at 5 years follow up (*p* = 0.016 relative to Cluster 4; Fig. 3D). Cluster 4 (blue) encompassed samples with a higher abundance of activated and exhausted memory CD4 and CD8 T cells, and activated and resting Tregs, whereas Cluster 2 (purple) had a low abundance of these T cell populations (Fig. 3C). Cluster 3 (yellow) was dominated by samples with a low abundance of B cells, and a high abundance of naive and memory CD4 and CD8 T cells. Patients with primary tumors of low grade were more frequent in this cluster. The clusters were significantly correlated to lymph node status, metastasis size and primary tumor histological grade, but were not correlated to BC subtype, later distant metastasis, tumor morphology, Ki-67 proliferation score, tumor size, invasiveness (invasive vs DCIS) of the primary tumor or the patient’s age (Fig. 3C).

### Modest differences in SN+ samples across subtypes

To investigate if BC subtypes had consequences for immune cell composition within lymph nodes, we compared the frequency of immune cell populations in SN- and SN+ separately. In SN- samples, no significant differences for individual immune cell subsets were observed between the different BC subtypes (Fig. 4A, Supplementary Table 3). However, we noted that for several cell populations, the frequency ranges were large, in particular within the ER^+^ subtype. In contrast to SN- samples, there was a significant increase in naive B cells between Her2^+^ and ER^+^ BC subtypes for SN+ samples. We also observed a trend towards an increase in GC B cells, plasma cells, and T follicular helper (T_FH_) cells in the TN subtype compared to in ER^+^ and Her2^+^ (Fig. 4B, Supplementary Table 3), suggesting a trend towards more active germinal center reaction in SN+ samples from TNBC patients.

**Figure 4:**
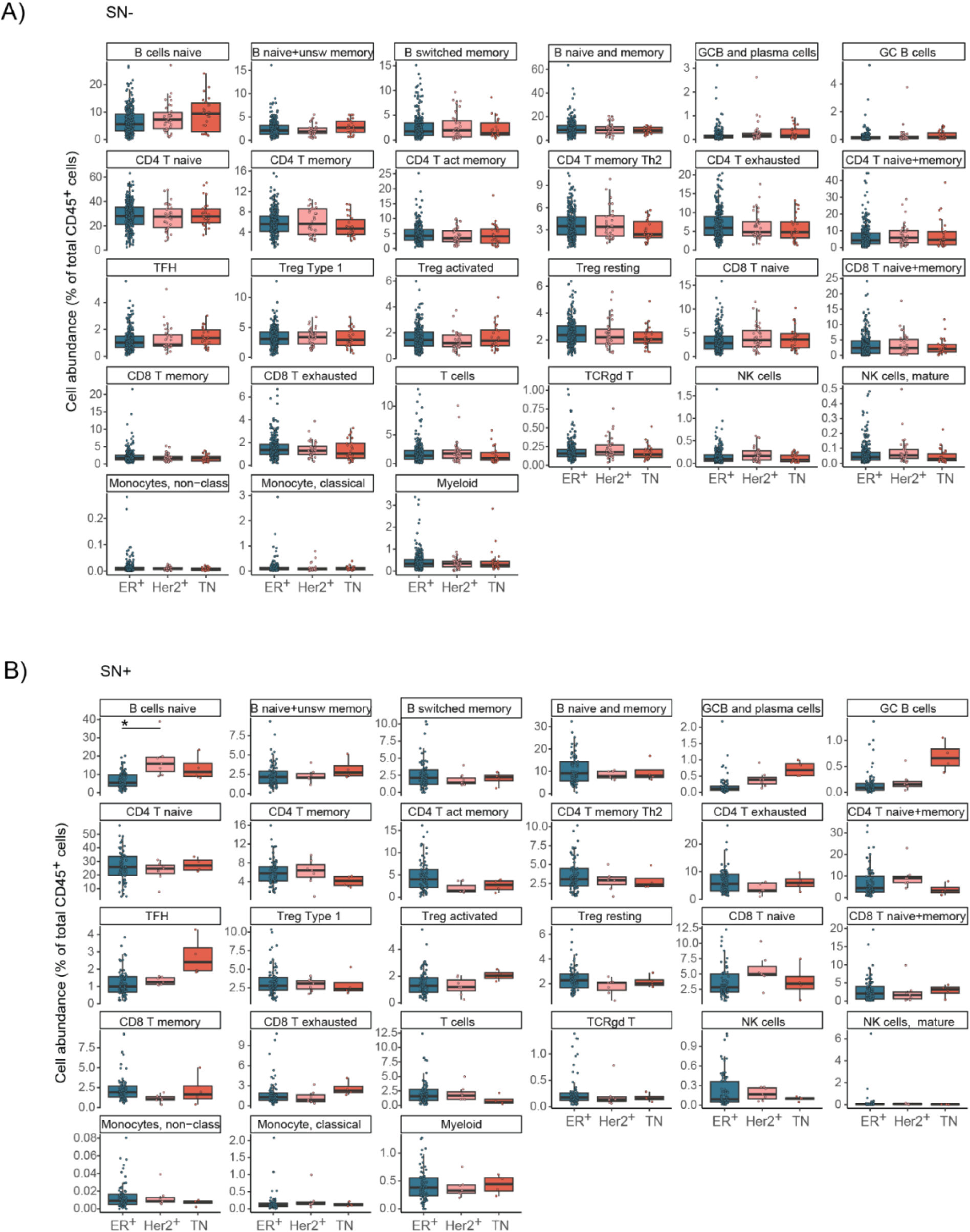
Immune cell composition in SN- and SN+ samples categorized by primary tumor subtypes. Immune cell frequencies in **A)** SN- and **B)** SN+ samples divided by the BC subtype of the primary tumors. Boxes indicate median value and upper and lower quartile, whiskers indicate maximum and minimum data values, and all sample values are shown as dots. \**p* < 0.05 (Mann-Whitney *U* test with Bonferroni correction (*n* = 27)).

### In ER^+^ patients, larger loci of metastatic cells had a greater impact on immune cell populations

ER^+^ breast cancer is the most prevalent subtype and is associated with increased lymph node metastasis (7,28), a pattern also reflected in this cohort. Finding a trend towards subtype-related differences in SN+ samples, we focused on samples from ER^+^ patients only, to address the immune response during progression within this subtype. The rate of distant metastasis for ALN+, SN+ and SN- samples from ER^+^ patients was similar to the whole cohort, likely due to the overrepresentation of ER^+^ patients, accounting for 79% of the cohort (Fig. 5A). ALN+ patients had significantly shorter DDFS compared to SN- and SN+ patients (*p* = 0.0000003 and *p* = 0.031, respectively), and SN+ had shorter DDFS than SN- patients (*p* = 0.0026).

**Figure 5:**
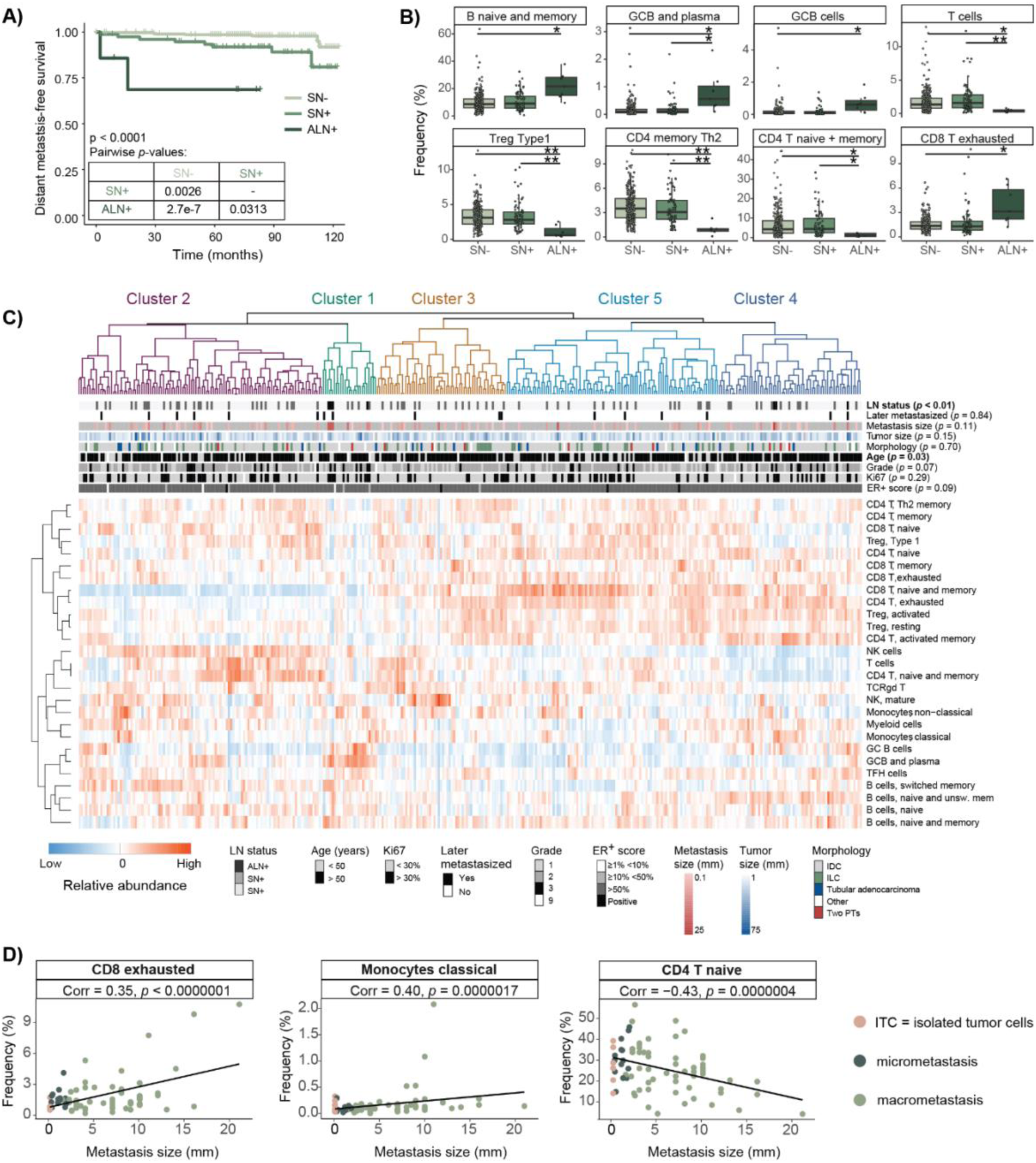
Differences in immune cell composition in lymph node samples from patients with ER^+^ primary tumors. A) Kaplan-Meier curves stratified by clinical lymph node status. Pairwise log-rank tests with Benjamini-Hochberg post-hoc correction were performed on the survival curves. **B)** Boxplot of selected immune cell populations of samples from ER^+^ patients (Mann-Whitney *U* test with Bonferroni correction (*n* =27), \**p*<0,05, \*\**p*<0,01). **C)** Hierarchical consensus clustering of cell type frequencies for all samples from ER^+^ patients. (*p*-values from Fisher’s exact test). Columns are normalized by total CD45+ cells per sample, and rows are normalized across all samples. Both are clustered by Pearson correlation distance matrix and average linkage. **D)** Correlation analysis of the abundance of selected immune cell populations vs. metastasis size for SN+ samples from ER^+^ patients (Spearman’s rank correlation). The straight line represents linear regression of the abundance of the immune cell population on the metastasis size. The dots are colored by sample type, corresponding to SN+ stratified by clinical metastasis size (ITC = isolated tumor cells). r_s_ = Spearman correlation coefficient.

When assessing the immune cell abundance across the lymph node stages, ALN+ samples had a significantly higher abundance of naive and memory B cells, GC B cells, plasma cells, and exhausted CD8 memory T cells, alongside a reduction of naive, memory and Th2 memory CD4 T cells, and Type 1 Tregs compared to SN-and SN+ samples (Figure 5B, Supplementary Fig. 5). No significant difference was found between SN- and SN+ samples. Hierarchical consensus clustering of samples from ER^+^ patients only revealed a small cluster significantly enriched in ALN+ samples and older patients (Cluster 1 (green)), encompassing relatively large metastases. This cluster had a particularly high abundance of GC B cells, plasma cells, and other B cells, and a low abundance of naive, memory and exhausted CD4 and CD8 T cells (Figure 5C). No significant correlation to other clinical parameters such as metastasis size, tumor size, later distant metastasis, tumor morphology, tumor grade or Ki-67 proliferation score was identified. Almost all samples were categorized with an estrogen score of >50%. No significant differences in DDFS were found when stratifying patients according to cluster identity (Supplementary Fig. 6).

As ALN+ samples harboring large metastases had a distinct immune cell composition compared to SN- and SN+ samples, we next investigated whether metastatic tumor burden within SN samples affected immune cell abundance. Linear regression of SN+ samples from ER^+^ patients revealed a moderate correlation between metastasis size and the abundance of naive CD4 T cells, classical monocytes and exhausted CD8 T cells. Specifically, naive CD4 T cells decreased as metastasis size increased, while classical monocytes and exhausted CD8 T cells increased (Fig. 5D, Supplementary Fig. 7), suggesting that the larger the loci of metastatic cells, the greater the impact on certain immune cell populations.

### SN- samples with detectable metastatic tumor cells had an immune profile similar to ALN+ samples

Since we observed a wide range in immune cell populations within SN- samples from ER^+^ patients and a large proportion of SN- samples clustered together with ALN+ samples, we performed separate analyses on SN- samples only. Studies on non-invasive (DCIS) and invasive primary tumors show increased immune infiltration in invasive tumors and DCIS with an invasive component compared to pure DCIS (29–31). This was not reflected in the SN from ER^+^ patients, where no significant differences in cell abundance between SN from invasive and non-invasive disease were found (Supplementary Fig. 8, Supplementary Table 4). This might suggest that the immune composition is altered already in a non-invasive setting, or that the changes are too small or local to be detected by this method.

As no significant differences were found between these samples, DCIS samples were included as SN- in further analyses. Comparing cell type abundance in SN-samples from ER^+^ patients stratified on clinical parameters revealed that patients over 50 years had a significantly lower abundance of naive CD8 T cells and significantly more activated Tregs compared to patients younger than 50 years. This observation aligns with previously reported findings of a decline in T cell populations and an increase in immunosuppressive cells with aging (32). No other significant differences associated with clinical parameters were observed (Supplementary Table 4). Due to the striking heterogeneity in the abundances of each immune cell population within SN- samples, we also calculated the abundance of tumor cells based on pan-cytokeratin positive (PanKer^+^) cells detected by CyTOF to reveal potential discrepancy with identification by histopathology. Tumor-cell positivity was defined as >0.02% tumor cells of total cell count in the sample. Seventy-three of the SN- samples from ER^+^ patients contained tumor cells, corresponding to 20.4% discrepancy with pathology classification (Supplementary Table 5).

Hierarchical consensus clustering revealed three distinct clusters (Fig. 6). Cluster 2 (purple) exhibited a suppressed immune profile, characterized by activated and resting Tregs, memory and exhausted CD8 T cells, and activated memory and exhausted CD4 T cells, which partly mimicked the immune profile of ALN+ samples. Interestingly, tumor cell identification by CyTOF was significantly correlated with the clustering and was enriched in cluster 2, which was characterized by an exhausted T cell phenotype (Fig. 6). In contrast, Cluster 1 (green) showed an increased abundance of B cells, including naive and switched memory B cells, GC B cells and plasma cells, NK cells and myeloid cells. Cluster 3 (yellow) was dominated by naive and memory CD4 T cells and Type1 T regulatory cells. No other significant correlations with clinical parameters were observed (Fig. 6). The clusters were, however, not related to differences in DDFS, but very few patients later experienced progression of the disease (*n* = 7) (Supplementary Fig. 9).

**Figure 6:**
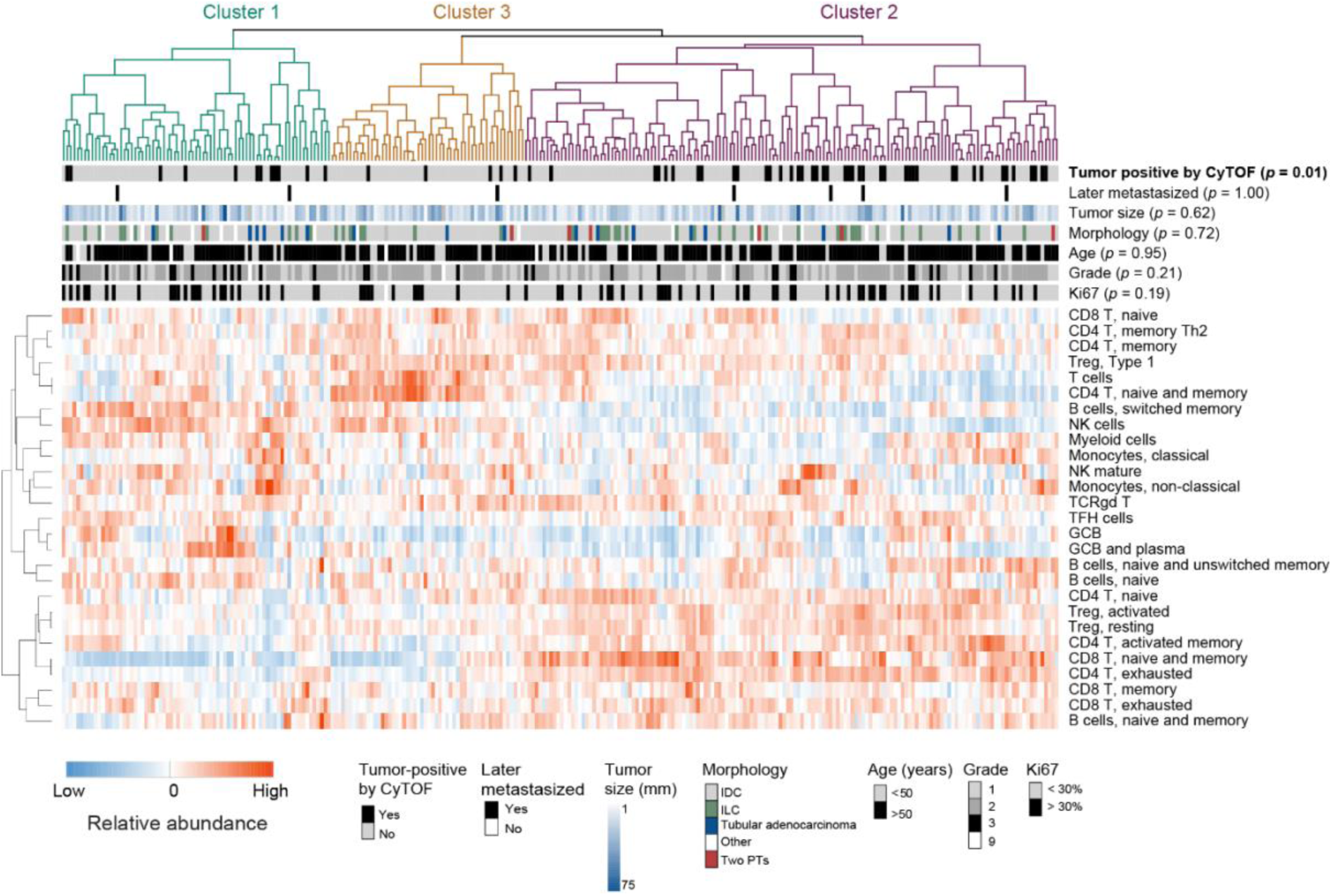
Hierarchical consensus clustering of SN- samples from ER^+^ patients (*p*-values from Fisher’s exact test). Columns are normalized by total CD45+ cells per sample, and rows are normalized across all samples. Both are clustered by Pearson correlation distance matrix and average linkage.

In total, our findings indicate that also smaller metastases can induce changes in the immune cell landscape towards T-cell exhaustion, and the degree of alterations are influenced by the metastatic size.

## Discussion

The prognostic significance of lymph node examination, particularly in axillary lymph nodes, is well-established, with larger lymph node metastases being strongly associated with a higher risk of distant metastasis (14,15). However, the clinical implications of smaller lymph node metastases remain a subject of ongoing debate. It has been reported that patients with smaller primary tumors (T1 and T2) and metastatic spread to the sentinel node rarely spread further (10,33). Other studies have shown that macro-metastases have worse prognosis compared to micro-metastases (34).

The role of lymph nodes in facilitating further dissemination also remains controversial. While some studies in breast cancer suggest limited seeding from lymph nodes to more distant organs (16), contrasting evidence from colon and prostate cancer suggests a potential link between lymph node metastases and the origin of distant metastases (17,18). Further complicating the picture is the observed heterogeneity within lymph node metastases. Studies have shown that a single lymph node can harbor multiple subclones, raising questions about the metastatic potential of these subpopulations (35). However, how tumor cells potentially alter the immune cell composition in lymph nodes during progression and whether there are BC subtype differences remains unclear. Here, we performed deep single-cell profiling of immune cells from lymph nodes with or without metastases, across 458 treatment-naive patients with ER^+^, Her2^+^ or TN breast cancer, encompassing 45 million single cells.

Our cohort has a relatively long follow-up time of 10 years, and the ER^+^ subtype was the dominant subtype in our study, accounting for 79% of cases, hence reflecting the real-world subtype composition of BC. In the total cohort, as well as the ER^+^ subcohort, ALN+ cases had the shortest time to distant metastasis, followed by SN+. Due to few ALN+ samples, these results should however be interpreted with caution. When regarding BC subtypes, it is known that ER^+^ breast cancers have a higher degree of lymph node involvement than TNBC, which tend to metastasize to distant organs independent of lymph node metastases (7). Despite no significant difference between the subtypes at 10 years follow-up, a significant difference was observed at 5 years follow-up where patients with TNBC performed worse than patients with ER^+^ and Her2^+^ disease. This is in line with the knowledge that TNBC has the highest 5-year recurrence rate (36,37).

ALN+ samples across all BC subtypes had a significant increase in exhausted CD8 T cells and memory CD8 T cells, and a decrease in naive CD4 T cells and resting Tregs as compared to SN- cases, in line with our prior results (24,25). Similar changes were found for ALN+ samples from ER^+^ cases. It has been suggested that Tregs stem from antigen-experienced effector CD4 T cells converted into induced Tregs in the tumor microenvironment (38). In addition to these T cell subset alterations, ALN+ samples from ER^+^ patients had notable alterations characterized by an increased abundance of naive and memory B cells and germinal center (GC) B cells, suggesting a modest activation of germinal center reactions. SN+ samples from TN patients also had a trend towards an increase in cell subsets related to a germinal center reaction, including GC B cells, plasma cells, and follicular helper T (T_FH_) cells as compared to Her2^+^ and ER^+^ samples. This suggests that smaller TNBC metastases in the sentinel node provoke an immune response similar to that seen in more advanced ALN+ metastasis from ER^+^ tumors. Whether this phenomenon is driven by the higher mutational burden typically observed in TNBC, or other subtype-specific factors or differences in tumor progression remains to be determined. TNBC is known for its increased immune infiltration at the primary tumor (4), and prior studies have reported a higher frequency of germinal centers in cancer-free lymph nodes of TNBC patients compared to non-TNBC cases (39). This suggests that TNBC cells are more immunogenic. Furthermore, the presence of germinal centers in lymph nodes have been associated with a lower risk of distant metastasis in TNBC, regardless of the amount of tumor-infiltrating lymphocytes at the primary tumor site (40). Our analysis, however, did not reveal any difference in immune cell abundance in SN- samples across subtypes. This could be explained by high variation in abundance for the different immune cell populations or differences in methodology. While the presence of germinal centers in non-involved lymph nodes is associated with improved survival in TNBC, studies in mice have reported that the primary tumor induced B cell accumulation in the draining lymph nodes (41). These B cells selectively promoted lymph node metastasis by producing pathogenic IgG which promotes metastasis, thus suggesting that B cells in tumor-draining lymph nodes might also play a tumor-promoting role.

SN- samples accounted for 74% of all lymph nodes in our cohort, so an important question to address was whether we could detect early changes in the immune cell composition prior to tumor cell invasion. We did not observe any difference in immune cell subset abundance in SN- samples across subtypes. However, when including only SN- samples from ER^+^ BC, unsupervised hierarchical cluster analysis revealed a cluster characterized by higher levels of activated and resting Tregs, and higher levels of memory and exhausted CD4 and CD8 T cells, partly similar to changes in ALN+ samples. This cluster was enriched for SN- samples with detectable levels of tumor cells by mass cytometry, despite being classified as SN- by pathology examination. Since epithelial cells are often underestimated in single-cell suspensions, the actual number of tumor positive cases are likely higher. Additionally, we did not analyze the same part of the lymph node as the pathologist, as each lymph node was cut into two halves for the two types of analysis. It is therefore likely that the part for pathological examinations lacked tumor cells. The presence of tumor cells in these samples suggests immunosuppression and T-cell exhaustion driven by the metastatic tumor cells. The cluster enriched in samples categorized as tumor-positive was however not correlated to later metastases (DDFS). Since ER^+^ patients commonly metastasize after 10 years (42,43), the follow-up data in this cohort is inconsistent and do not extend much beyond that period. The number of patients suffering from distant metastasis is likely to increase in the years to come. Prior studies have also detected increased Treg/CD4 T cell ratio in SN+ samples, but also demonstrated that this ratio was increased in SN+ samples as compared to ALN from women undergoing prophylactic mastectomy (44). A gradual increase in myeloid derived suppressor cells (MDSC) was found in SN- and SN+ as compared to disease-free ALN samples (44).

While lymph nodes may not be the primary source of distant metastasis, their involvement often reflects the metastatic potential of the tumor cells. The ability of tumor cells to colonize and alter the lymph node microenvironment suggests a similar capacity to invade and metastasize to other organs. Understanding the microenvironmental alterations, including subtype-specific variations, induced by tumor cells within the lymph node can offer valuable insights into the biology of the primary tumor and its metastatic potential. This knowledge can aid in identifying potential therapeutic targets.

The ER^+^ subtype was the dominant BC subtype in our study and is representative of the real world of BC. As our Her2+ and TNBC sub-cohorts were much smaller, comparisons across subtypes might be underpowered. Further limitations of our study include a lack of lymph nodes from healthy donors as controls for SN- samples. This is work in progress, but access to healthy lymph nodes may be an ethical challenge. As our mass cytometry analysis is based on single-cell suspensions, minor local alternations in SN might not be detected, as we have demonstrated in a case study (25). Tracking of tumor-specific alterations requires single-cell RNA-seq and TCR-seq of both primary tumors and lymph nodes. Recent studies of paired primary tumor and metastatic lymph nodes have illuminated metastatic seeding of genetically distinct clones (35,45), and the team of Yates demonstrated that distinct genetic metastatic clones could occupy different immune cell microenvironments within the same LN (35). Lastly, the subtyping of our samples is done by immunohistochemical staining, however molecular subtyping using PAM50 will be performed and may provide more distinct changes between subtypes.

This is the largest study characterizing the immune profile in lymph nodes of breast cancer patients, offering insight into the changes in the immune microenvironment during breast cancer progression. Our study underscores disease stage and tumor burden as the most relevant factors for immune suppression in the axillary lymph nodes. Indications of early immune/GC activation in SN+ samples were found for smaller TNBC metastases, an alteration occurring in the more manifested ALN+ samples. Our identification of an ER+ subgroup with distinct immune activation and suppression categorized with clinically tumor-negative sentinel nodes, but tumor-positive according to mass cytometry, enables further stratification of patients which may be biologically relevant. By accounting for the normal variation in immune cell populations within normal lymph nodes, tumor-specific alterations may be determined, thus enabling the identification of immune profiles predictive for therapeutic responses or prognosis.

## Consortia

### Oslo Breast Cancer Consortium (OSBREAC)

Kristine Kleivi Sahlberg, Elin Borgen, Anne-Lise Børresen-Dale, Olav Engebråten, Britt Fritzman, Øystein Garred, Jürgen Geisler, Gry Aarum Geitvik, Solveig Hofvind, Vessela N Kristensen, Rolf Kåresen, Anita Langerød, Ole Christian Lingjærde, Gunhild Mari Mælandsmo, Hege G Russnes, Torill Sauer, Helle Kristine Skjerven, Ellen Schlichting, Therese Sørlie

## Data availability

Data will be deposited in a suitable repository prior to publication.

## Acknowledgements

We thank Eldri Undlien Due and Inger Riise Bergheim for support in biobanking samples. This work was supported by grant from South-Eastern Norway Regional Health Authority (grant 2019045).

## Ethics approval and consent to participate

The study was approved by the regional committee for research ethics (200606181-1, 538-07278a, 2009/4935, 2016/433), and the samples were collected with written patient consent in accordance with the Declaration of Helsinki.

## Authors’ contributions

IHR and HR designed the study. MOF and KTL carried out experiments. MOF, KH, IHR and CL analyzed the data. ØG performed pathological evaluations. IHR acquired sample material. OCL supported biostatistical analysis. MLHR provided clinical support. MOF, JM, KH, IHR wrote the manuscript. IHR and KH supervised the study.

## Supplementary figures and tables

**Supplementary Figure 1:**
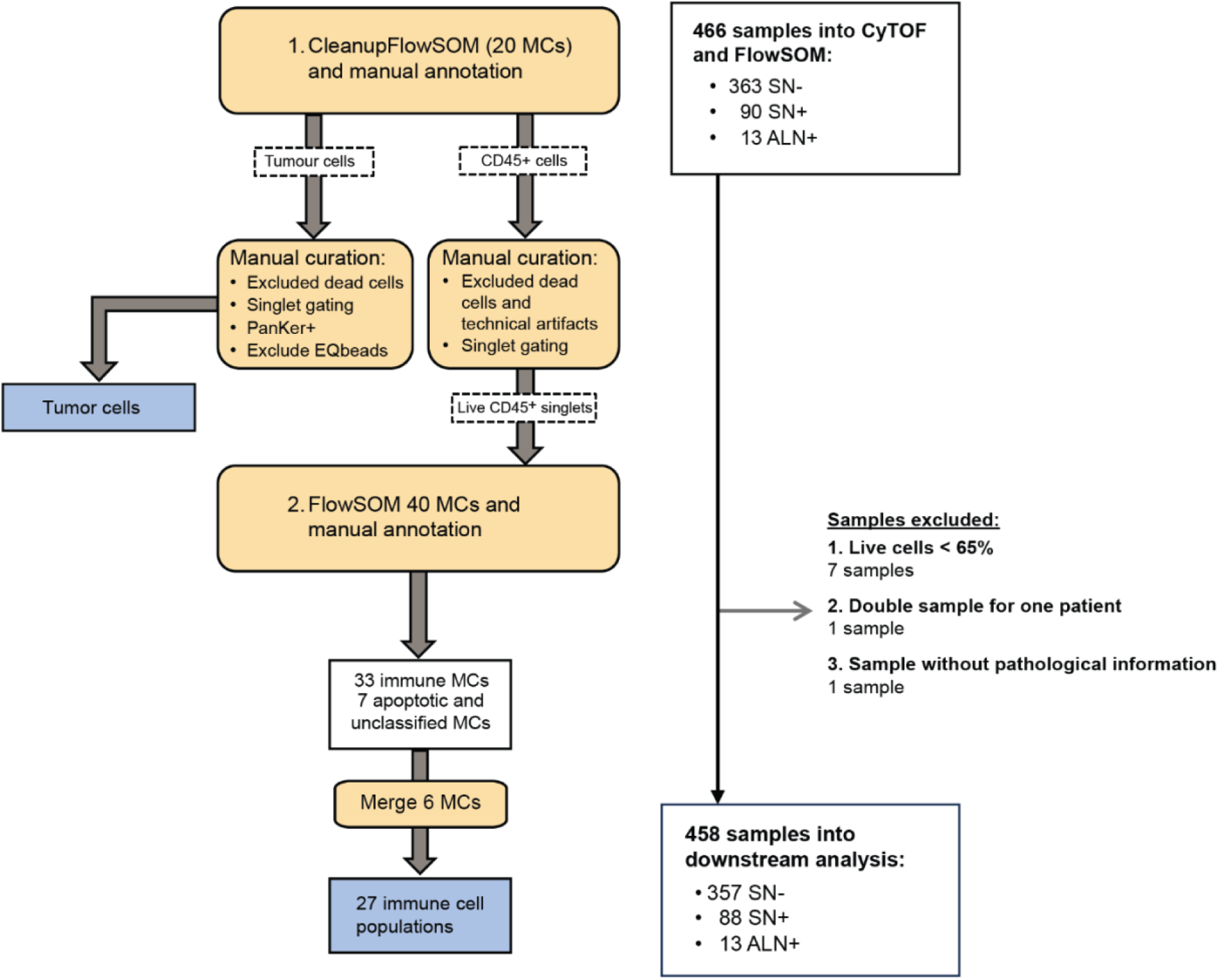
Flowchart of the sequential FlowSOM analysis strategy for semi-automatic gating and clustering of cells established in previous work (ref Fjørtoft), indicating included samples and the exclusion criteria. FlowSOM 1 was used for cleaning up samples and identification of CD45^+^ immune cells and PanKer^+^ tumor cells. Due to a high amount of tumor negative samples in the cohort, tumor cells were not included in the following FlowSOM 2, but manually curated before event counts were exported. FlowSOM 2 included 40 metaclusters (MCs), whereof 33 immune MCs, 3 apoptotic MCs and 4 former tumor cell MCs. The few cells residing in the tumor cell MCs were of different cell types and were together with the apoptotic MCs excluded from further analysis. 9 samples were excluded from the final analysis: 7 due to low viability, 1 due to two samples from the same patient, and 1 sample missing pathological lymph node classification. This resulted in 458 lymph node samples in the final analysis.

**Supplementary Figure 2:**
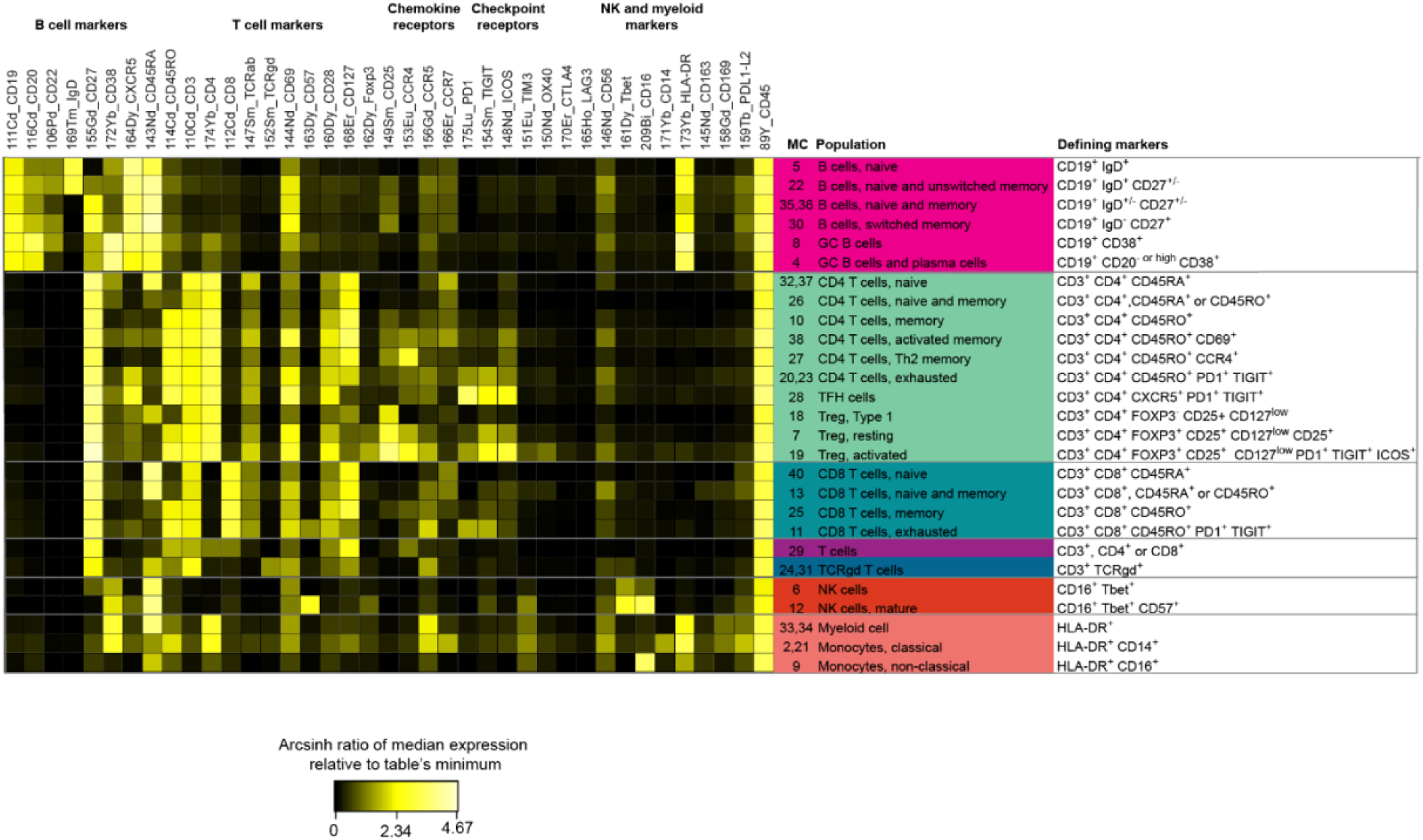
Merging and classification of the 33 MCs from FlowSOM2, resulting in 6 B cell populations, 16 T cell populations, 2 NK cell populations and 3 myeloid populations. The heatmap shows the arcsinh transformed ratio of the median channel expression relative to table’s minimum for 60 selected samples (289 000 cells) from all lymph node categories (SN- from non-invasive and invasive breast cancer, SN+ with ITC, micro-and macrometastasis, and ALN+ samples), concatenated using Cytobank FCS Concat Tool 0.7. Main cell markers driving the clustering are indicated. MCs were merged when they contained very similar cells.

**Supplementary Figure 3:**
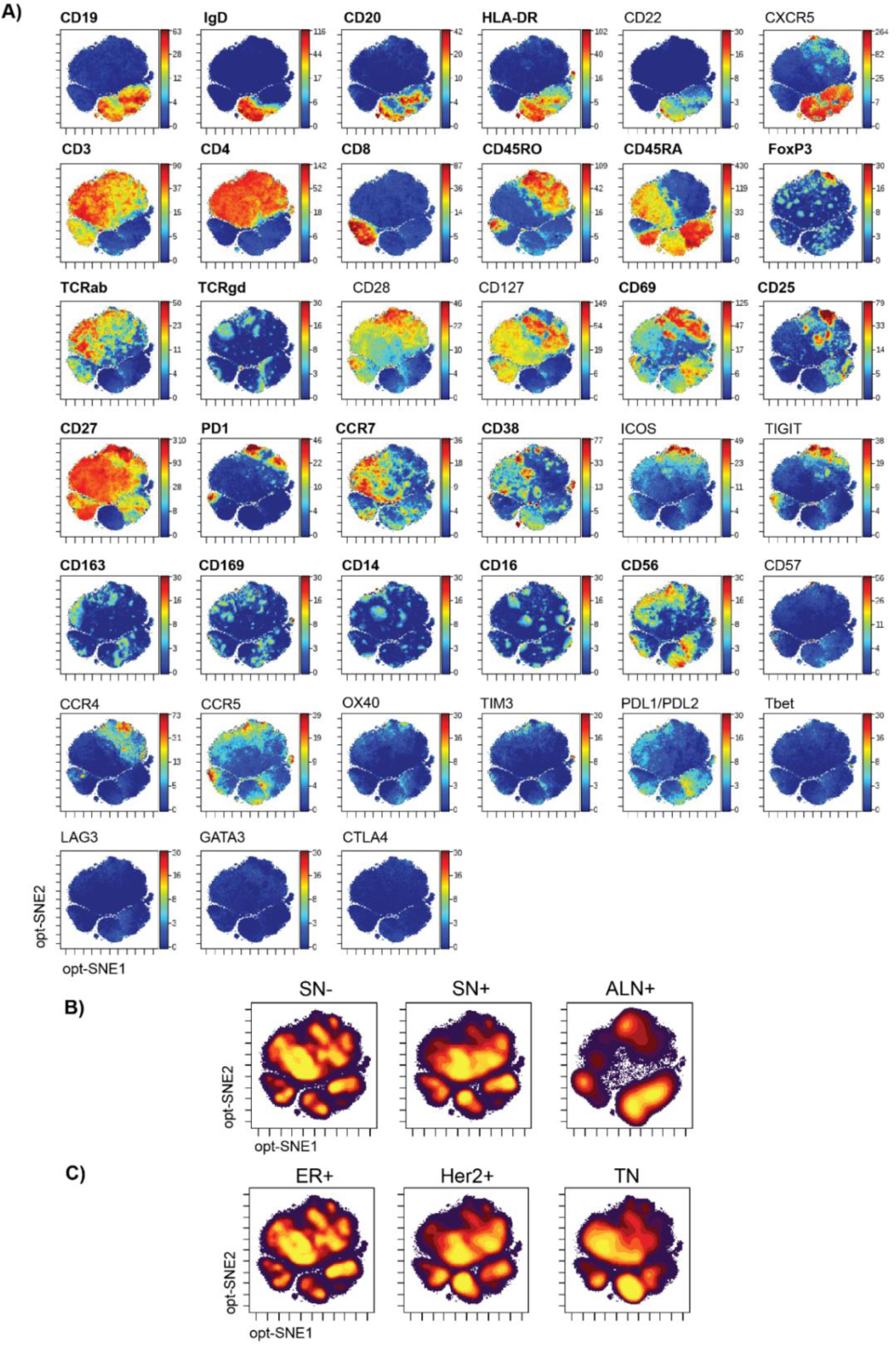
Expression of immune-cell markers on immune cells from all 458 lymph nodes, visualized by optSNE (markers in bold were included in the optSNE analysis). **A)** All samples concatenated and colored by markers as annotated. **B-C)** Density plot showing the distribution of immune cells across the different lymph node statuses **(B)** and in the three subtypes **(C)**. (SN- = sentinel node without metastasis, SN+ = metastatic sentinel node and ALN+ = metastatic axillary nodes).

**Supplementary Figure 4:**
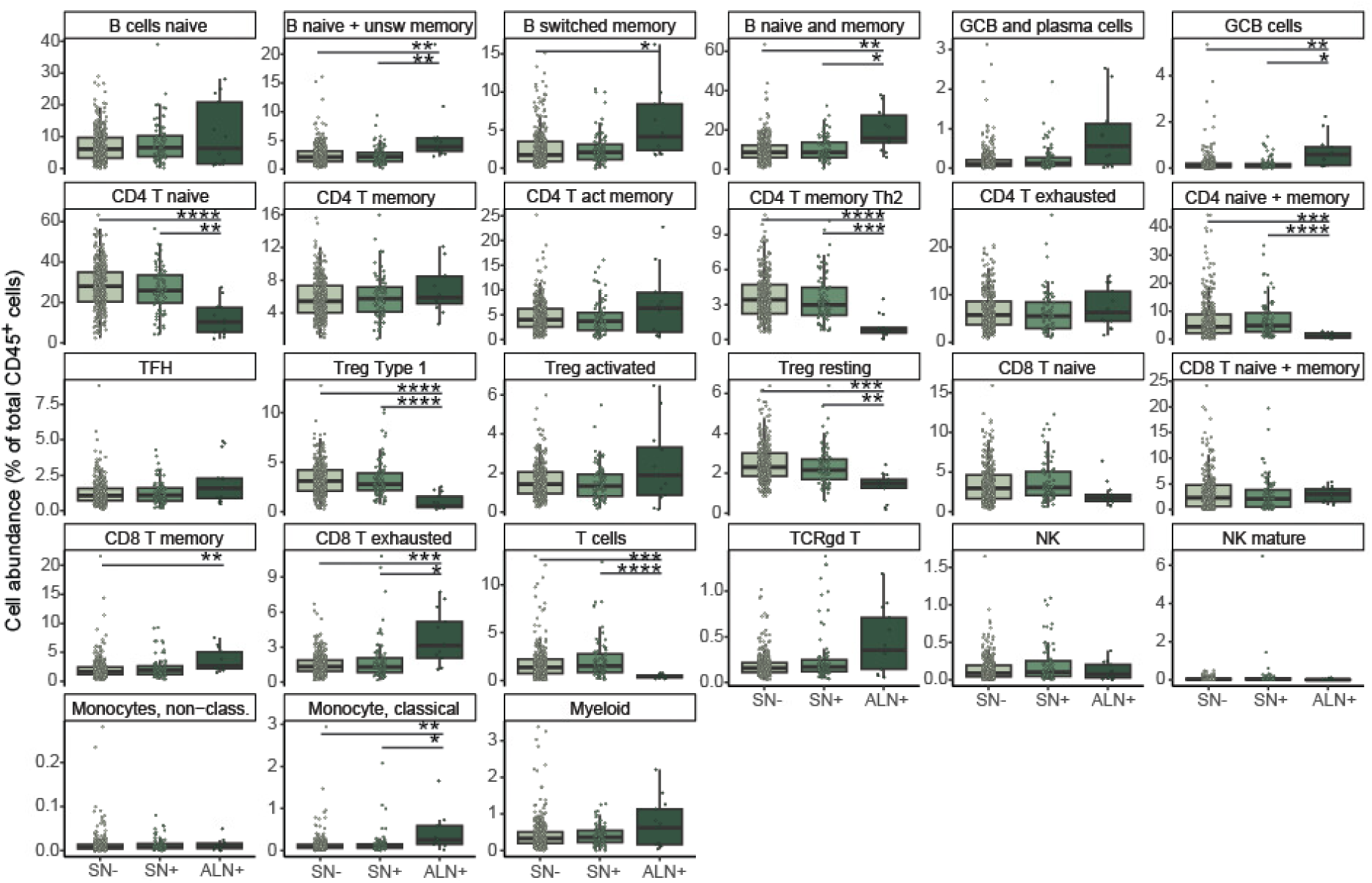
Differences in cell frequencies across tumor progression within all breast cancer subtypes (SN- = sentinel node without metastasis, SN+ = metastatic sentinel node and ALN+ = metastatic axillary nodes).

**Supplementary Fig. 5:**
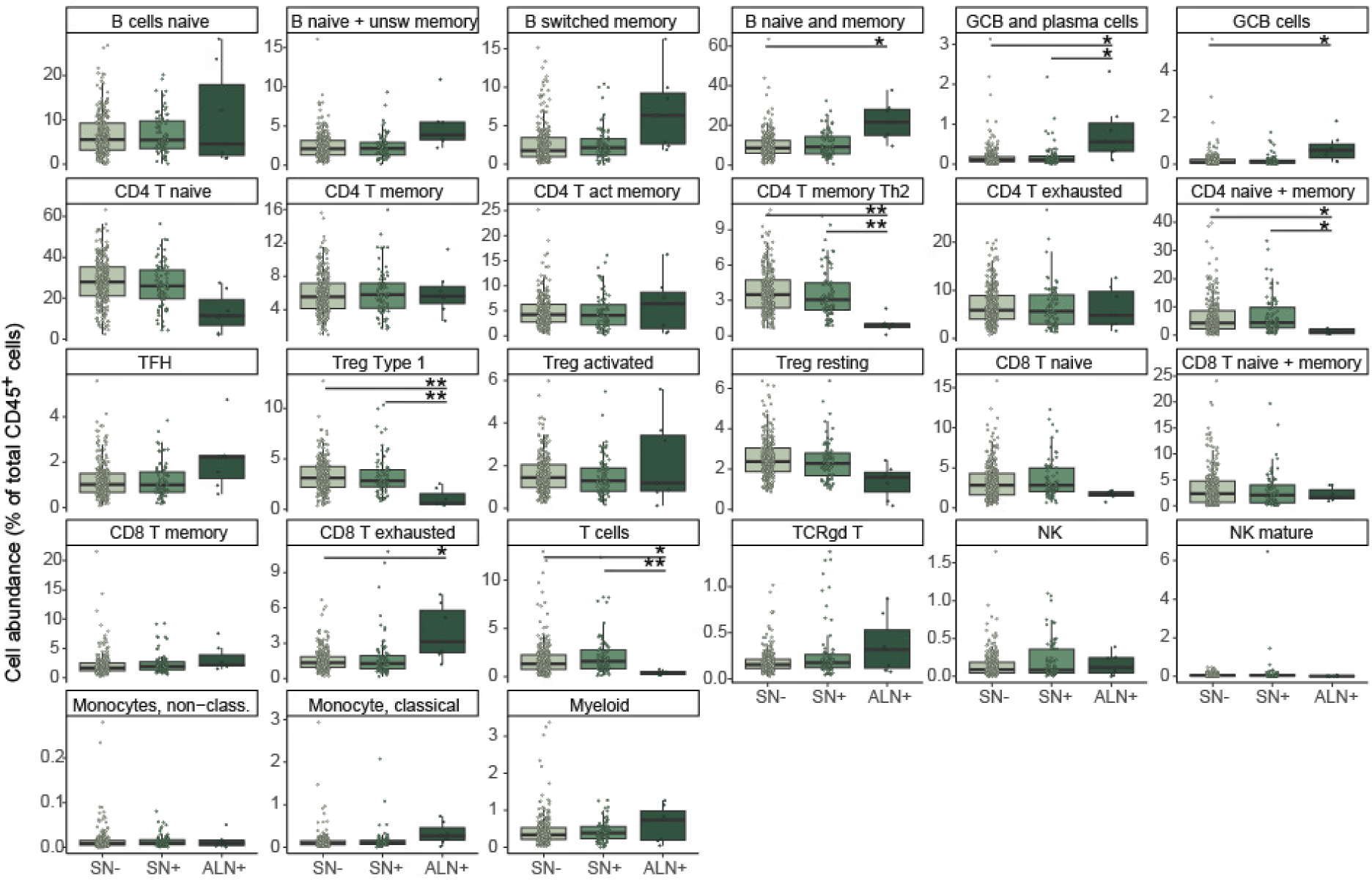
Differences in cell frequencies across tumor progression across all samples from ER+ patients (SN- = sentinel node without metastasis, SN+ = metastatic sentinel node and ALN+ = metastatic axillary nodes).

**Supplementary Figure 6:**
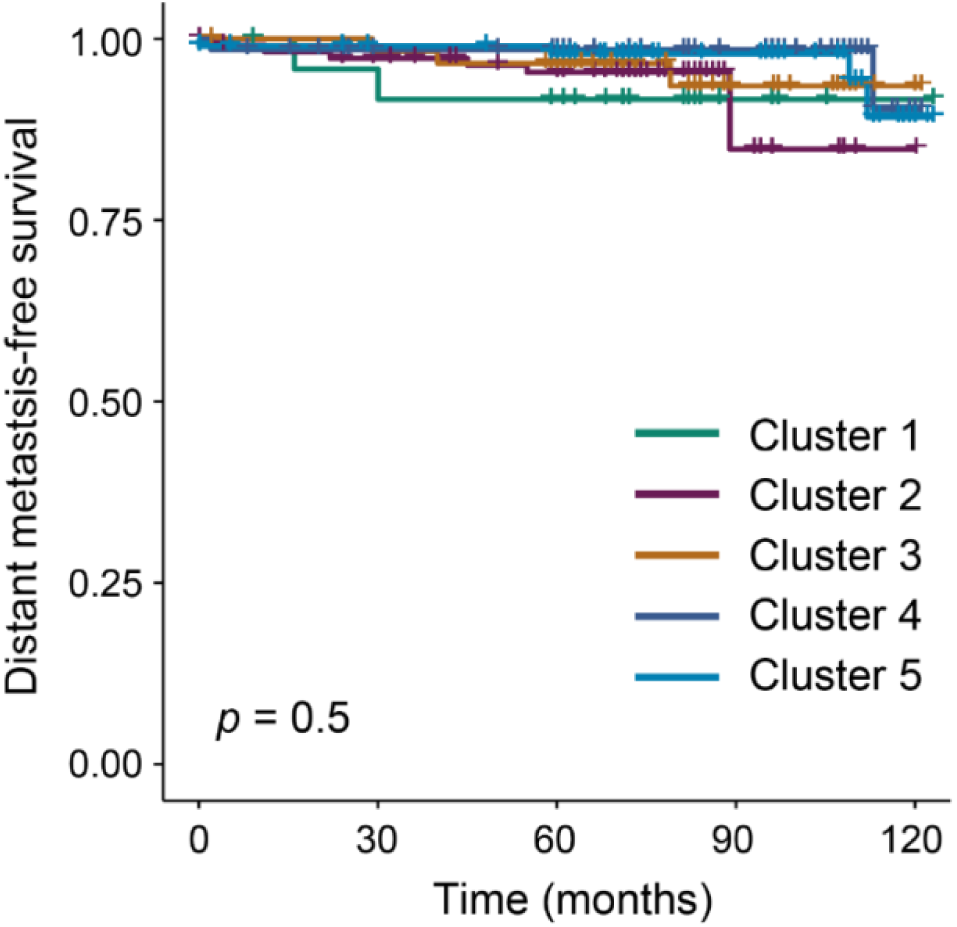
Kaplan-Meier curves showing disease-free survival (DFS) when stratifying patients according to cluster identity for ER+ samples only. Pairwise log-rank tests with Benjamini-Hochberg post-hoc correction was performed on the survival curves.

**Supplementary Fig. 7:**
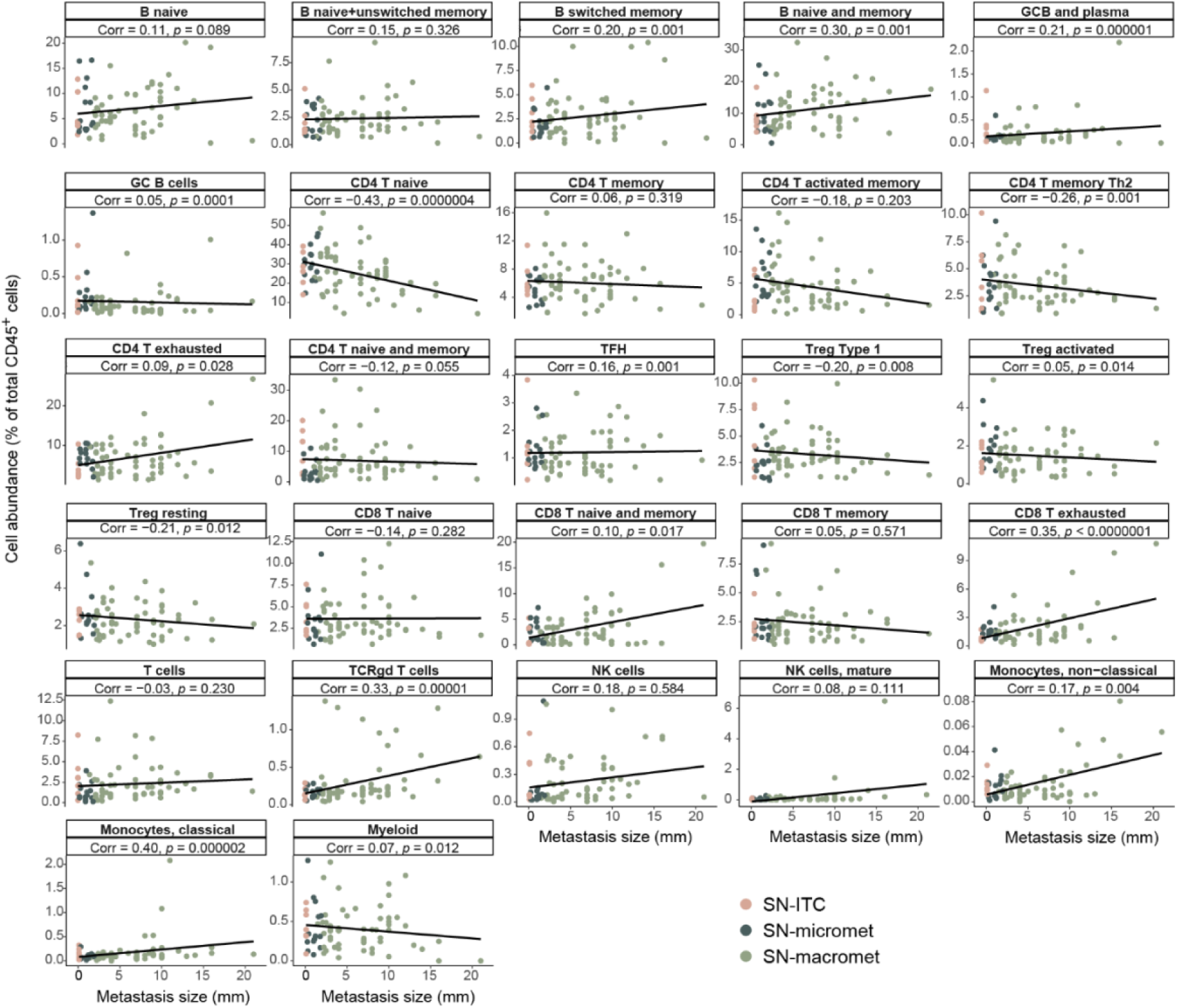
Linear regression of metastasis size vs. immune cell populations in SN+ samples. Samples are colored by metastasis size of SN+. (ITC = isolated tumor cells).

**Supplementary Fig. 8:**
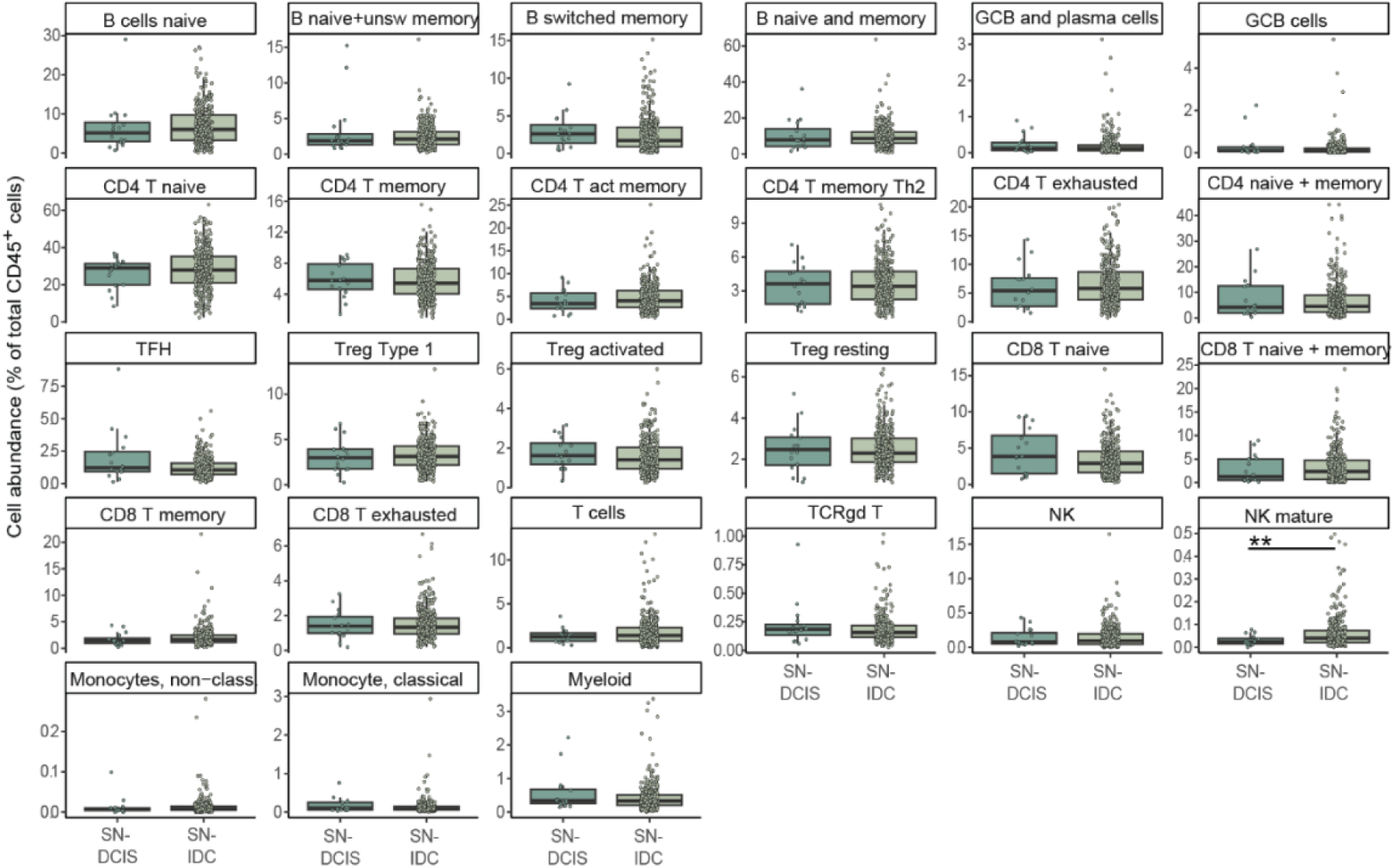
Differences between non-invasive and invasive primary tumors among SN- samples from ER+ patients. Immune cell frequencies of non-invasive DCIS samples and invasive SN- samples from ER+ patients. Boxes indicate median value and upper and lower quartile, whiskers indicate maximum and minimum data values, and all sample values are shown as dots.

**Supplementary Figure 9.**
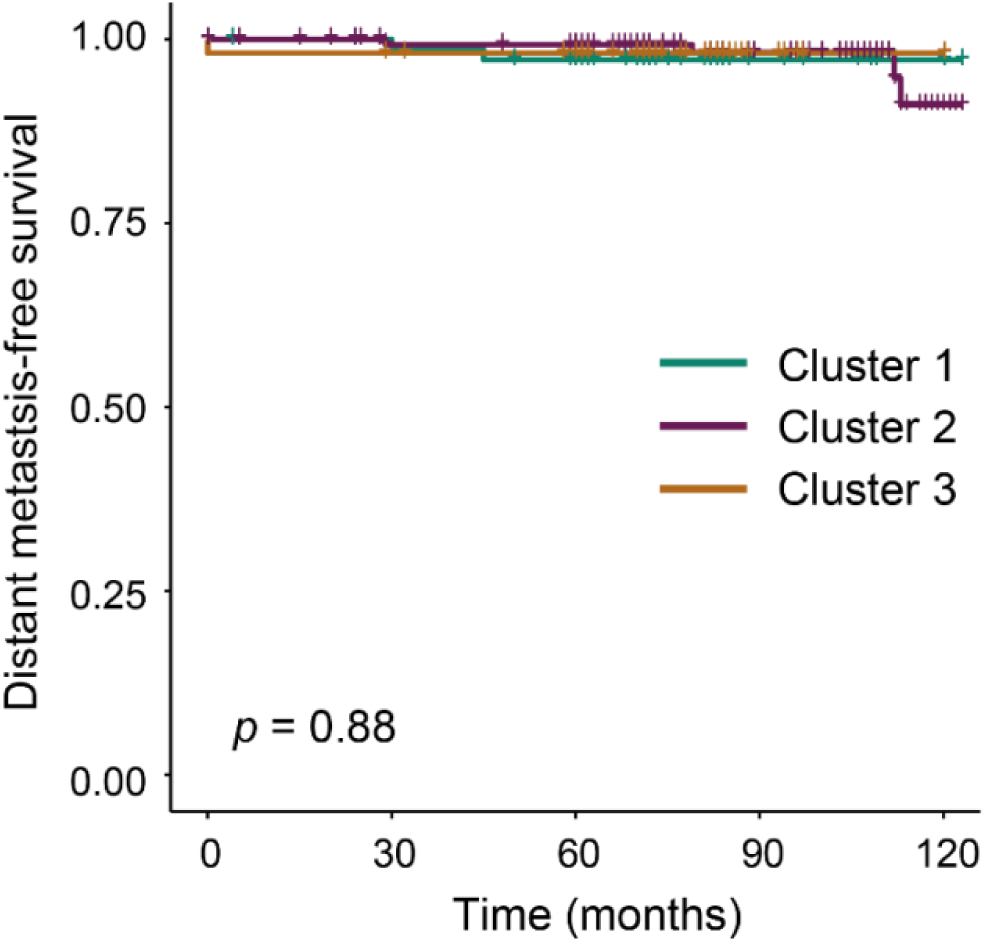
Kaplan-Meier curves showing disease-free survival (DFS) when stratifying patients according to cluster identity for SN- samples from ER+ patients only (Figure 6). Pairwise log-rank tests with Benjamini-Hochberg post-hoc correction was performed on the survival curves.

**Supplementary Table 1:**
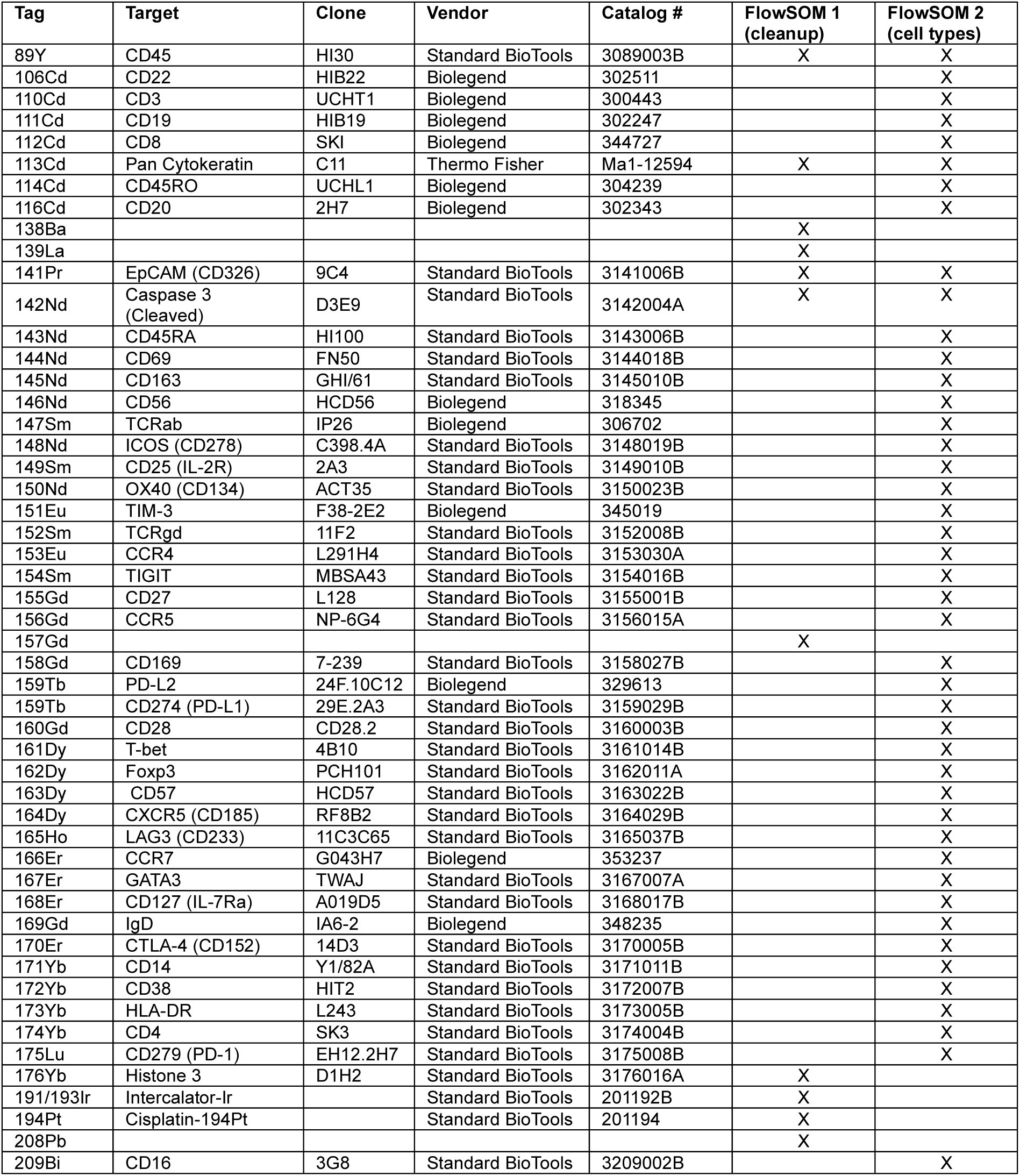
Antibodies and open channels in mass cytometry analysis and in FlowSOM analyses.

**Supplementary Table 2:**
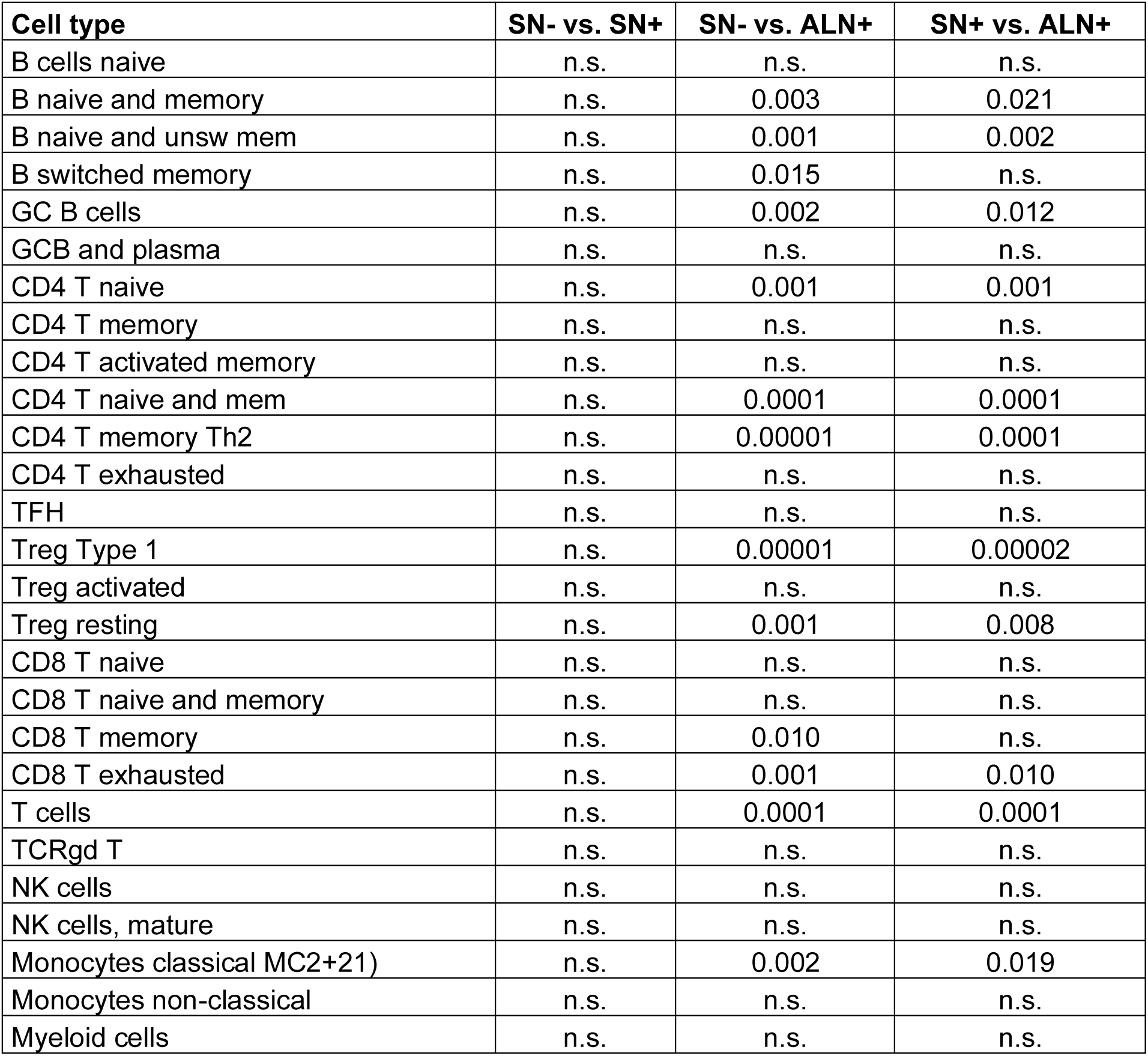
*p*-values from Mann-Whitney *U* test (after Bonferroni correction, *n* = 27) between all lymph node samples included in Figure 2 and Supplementary Fig. 4.

**Supplementary Table 3:**
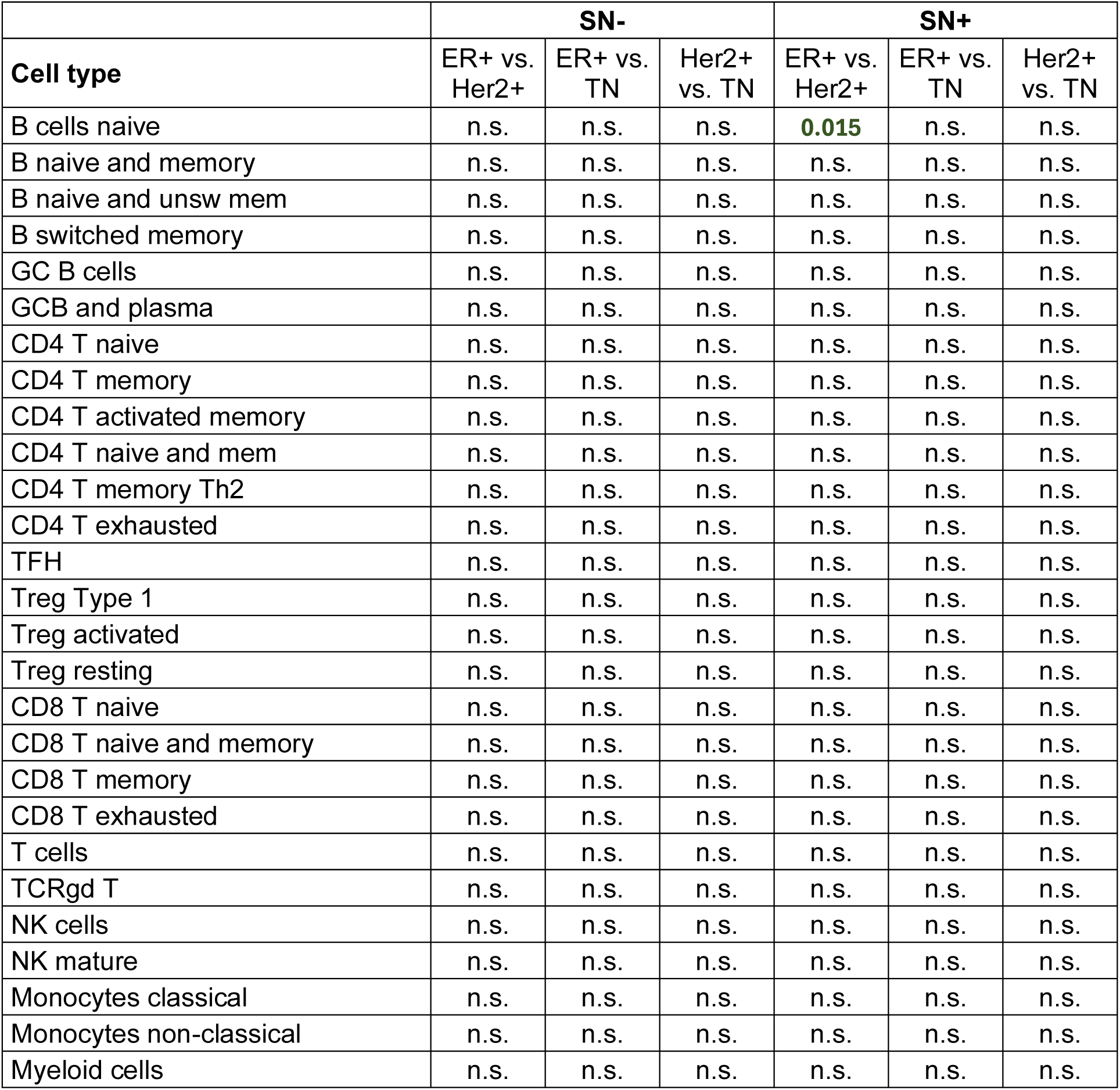
*p*-values from Mann-Whitney *U* test (after Bonferroni correction, *n* = 27) between breast cancer subtypes in SN- and SN+ samples, data included in Figure 4.

**Supplementary Table 3:**
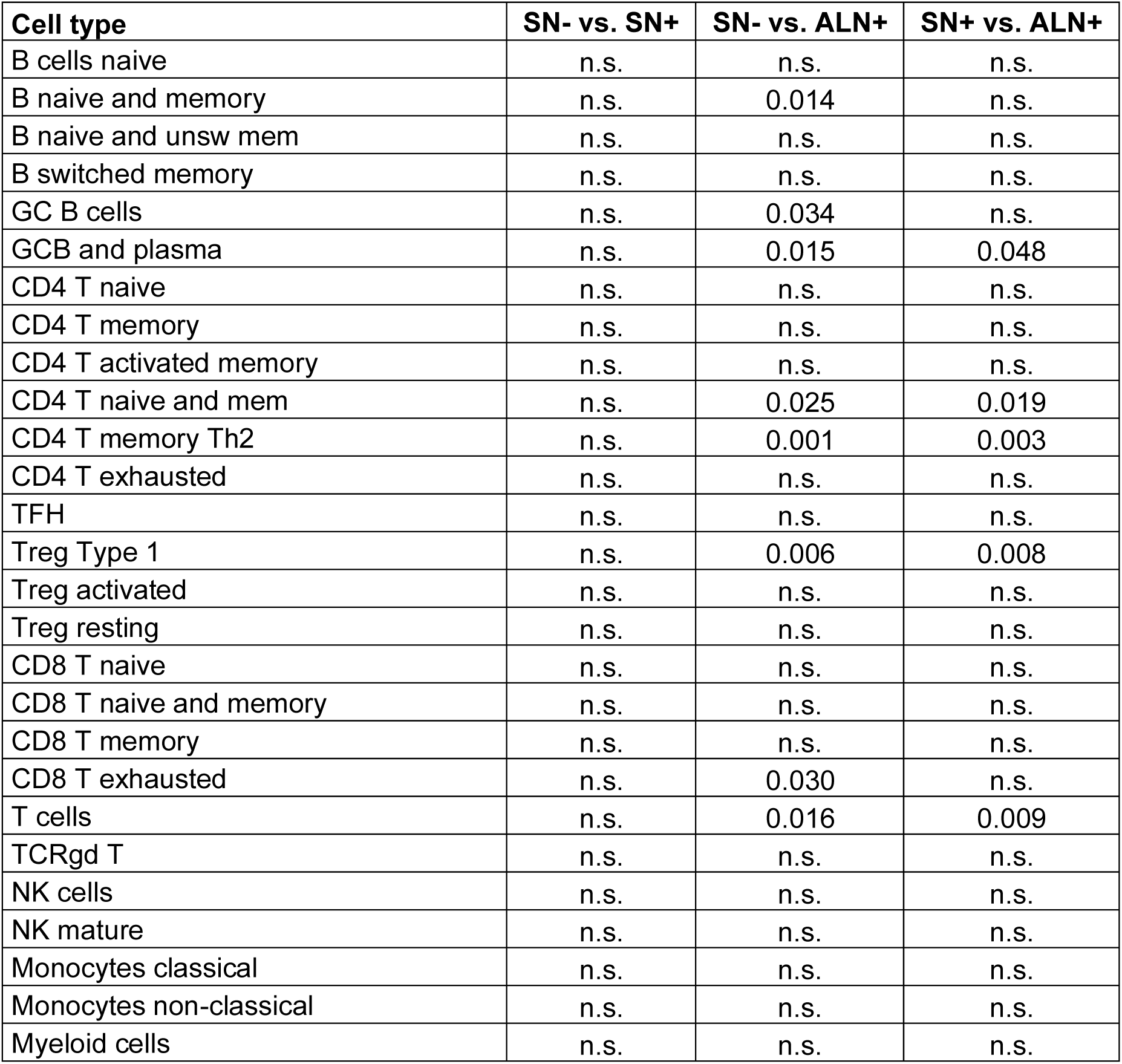
*p*-values from Mann-Whitney *U* test (after Bonferroni correction, *n* = 27) between breast cancer subtypes in lymph node samples from ER+ patients included in Figure 5 and Supplementary Fig. 5.

**Supplementary Table 4:**
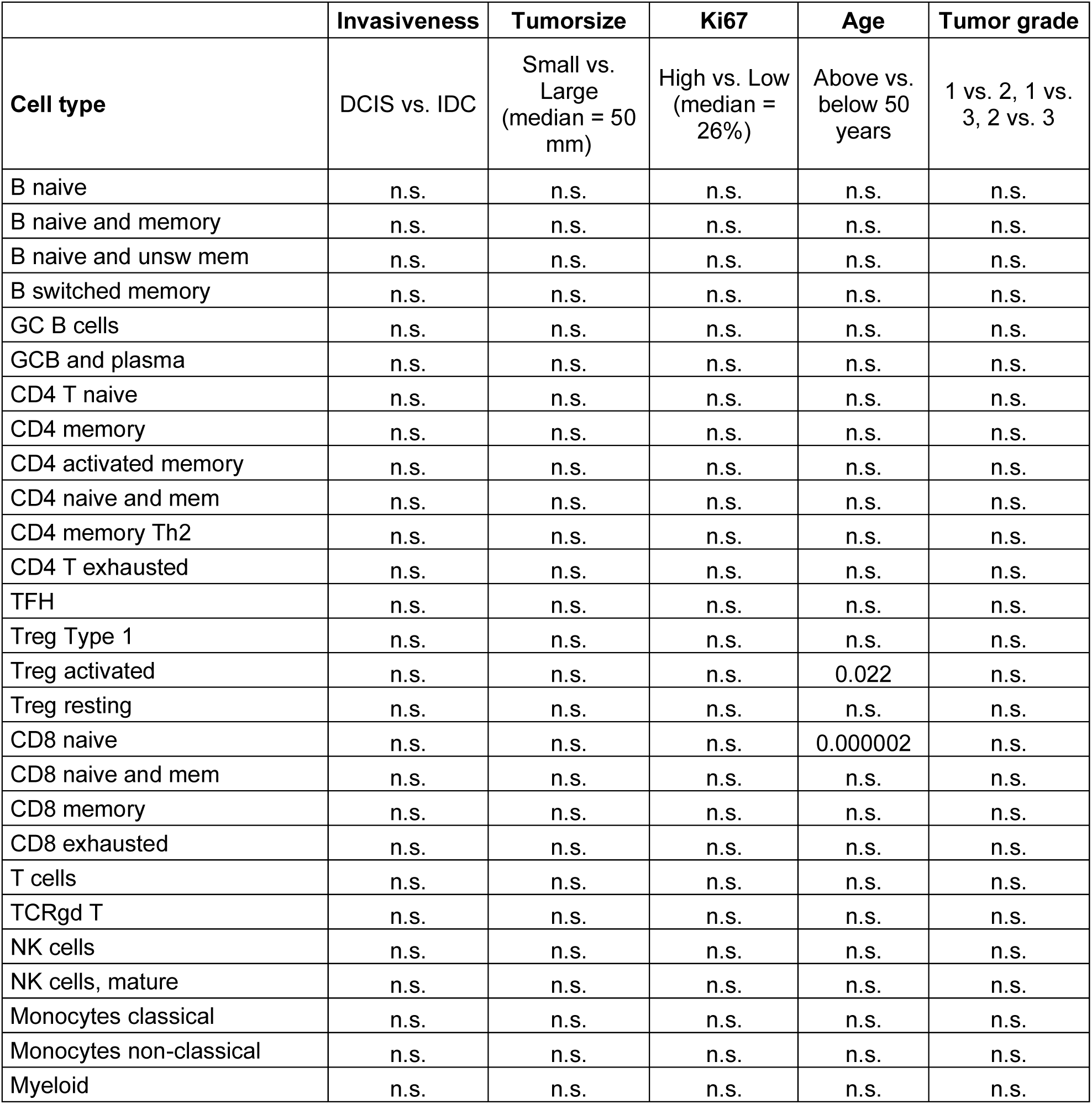
*p*-values from Mann-Whitney *U* test (after Bonferroni correction, *n* = 27) between different clinical parameters in SN- samples from ER+ patients.

**Supplementary Table 5:**
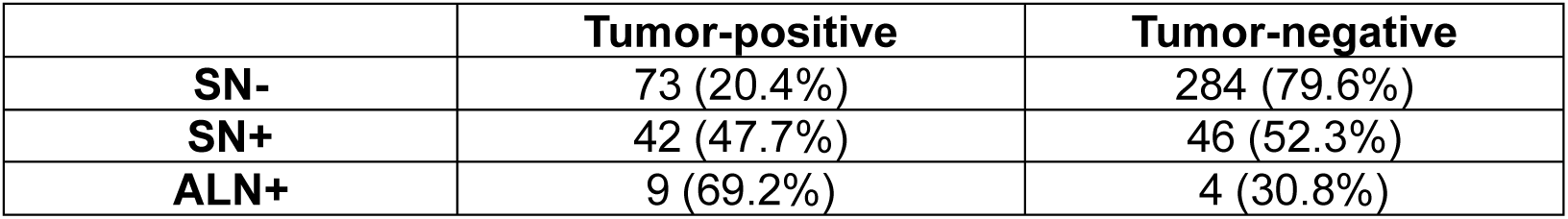
Number of samples with tumor cells detected by mass cytometry (CyTOF) analysis in SN-, SN+ and ALN+ lymph nodes. Samples were defined as tumor-positive when tumor cells were >0.02% of total cells.

