## Supplemental files for "Single-cell immune profiling of regional lymph nodes during early-stage breast cancer progression"

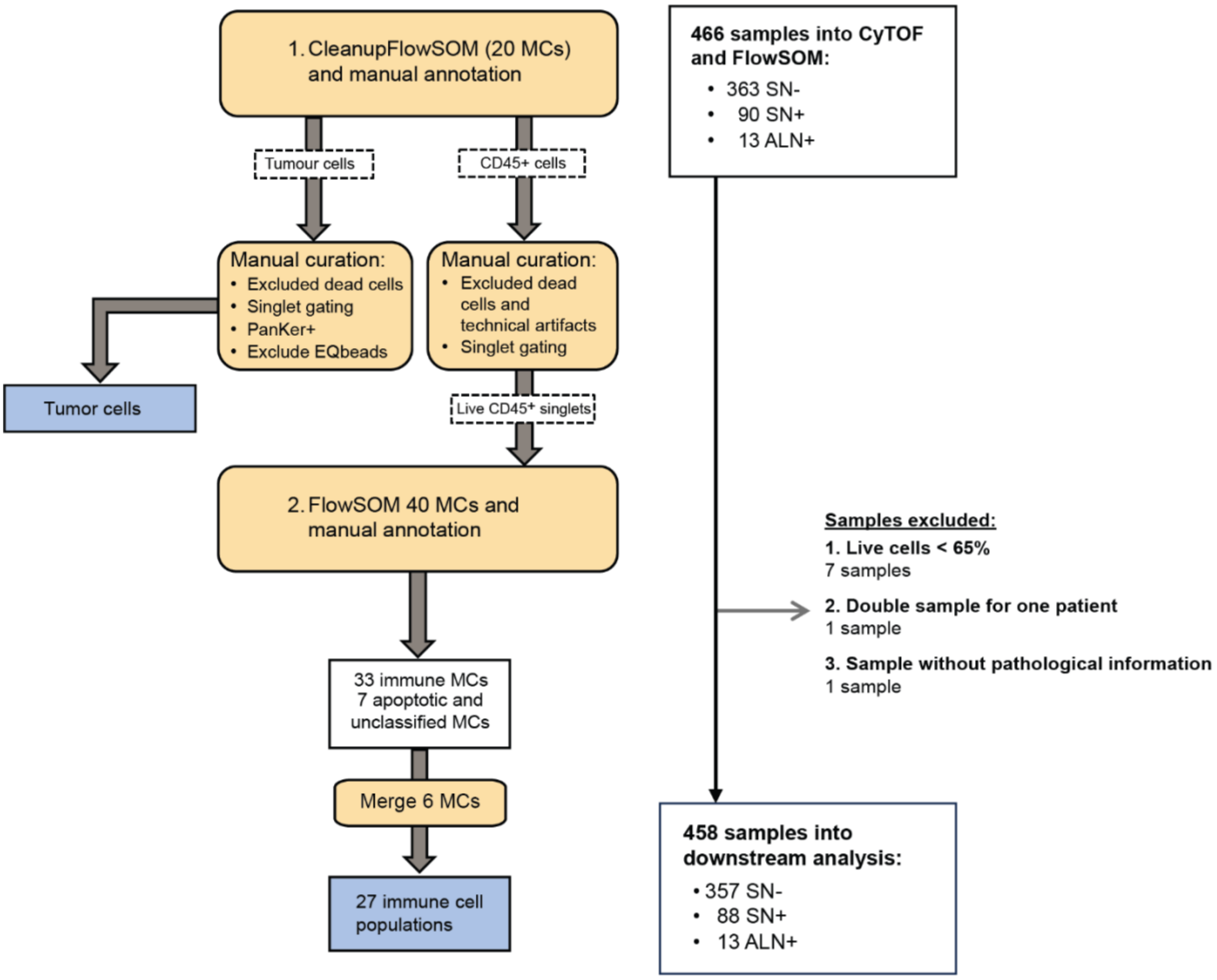


**Supplementary Figure 1:** Flowchart of the sequential FlowSOM analysis strategy for semi-automatic gating and clustering of cells established in previous work (ref Fjørtoft), indicating included samples and the exclusion criteria. FlowSOM 1 was used for cleaning up samples and identification of $CD45^+$ immune cells and $PanKer^+$ tumor cells. Due to a high amount of tumor negative samples in the cohort, tumor cells were not included in the following FlowSOM 2, but manually curated before event counts were exported. FlowSOM 2 included 40 metaclusters (MCs), whereof 33 immune MCs, 3 apoptotic MCs and 4 former tumor cell MCs. The few cells residing in the tumor cell MCs were of different cell types and were together with the apoptotic MCs excluded from further analysis. 9 samples were excluded from the final analysis: 7 due to low viability, 1 due to two samples from the same patient, and 1 sample missing pathological lymph node classification. This resulted in 458 lymph node samples in the final analysis.

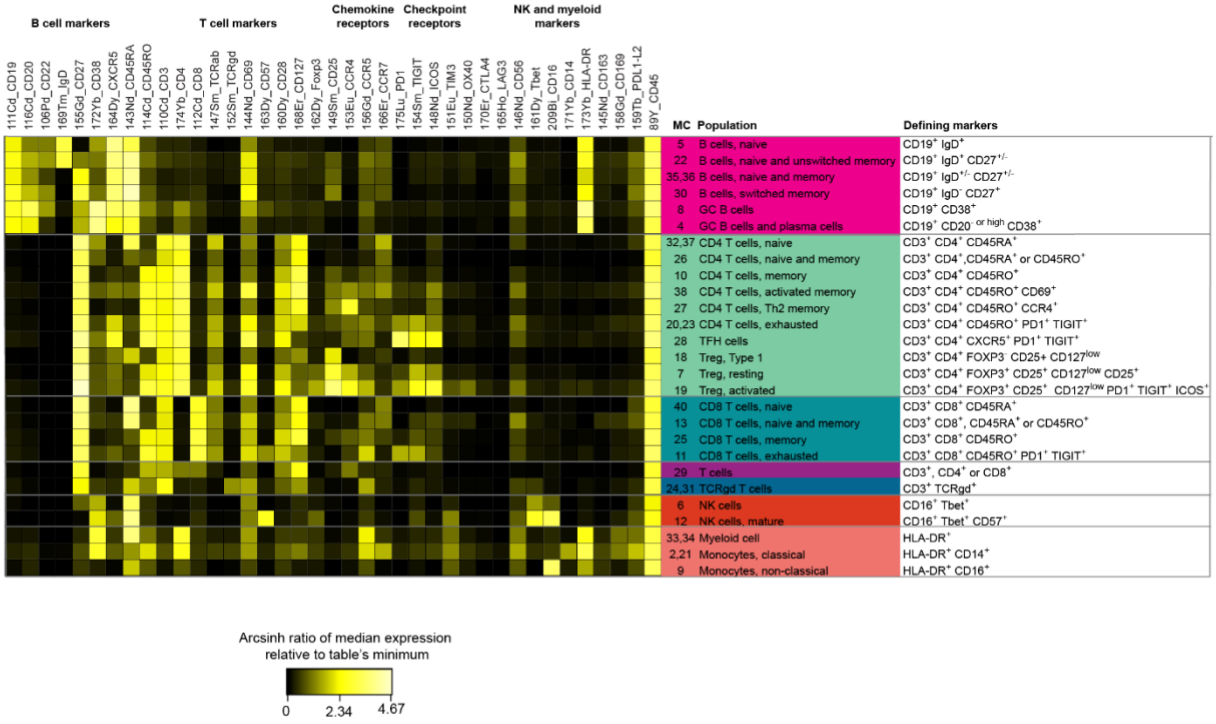


**Supplementary Figure 2:** Merging and classification of the 33 MCs from FlowSOM2, resulting in 6 B cell populations, 16 T cell populations, 2 NK cell populations and 3 myeloid populations. The heatmap shows the arcsinh transformed ratio of the median channel expression relative to table's minimum for 60 selected samples (289 000 cells) from all lymph node categories (SN- from non-invasive and invasive breast cancer, SN+ with ITC, micro- and macrometastasis, and ALN+ samples), concatenated using Cytobank FCS Concat Tool 0.7. Main cell markers driving the clustering are indicated. MCs were merged when they contained very similar cells.

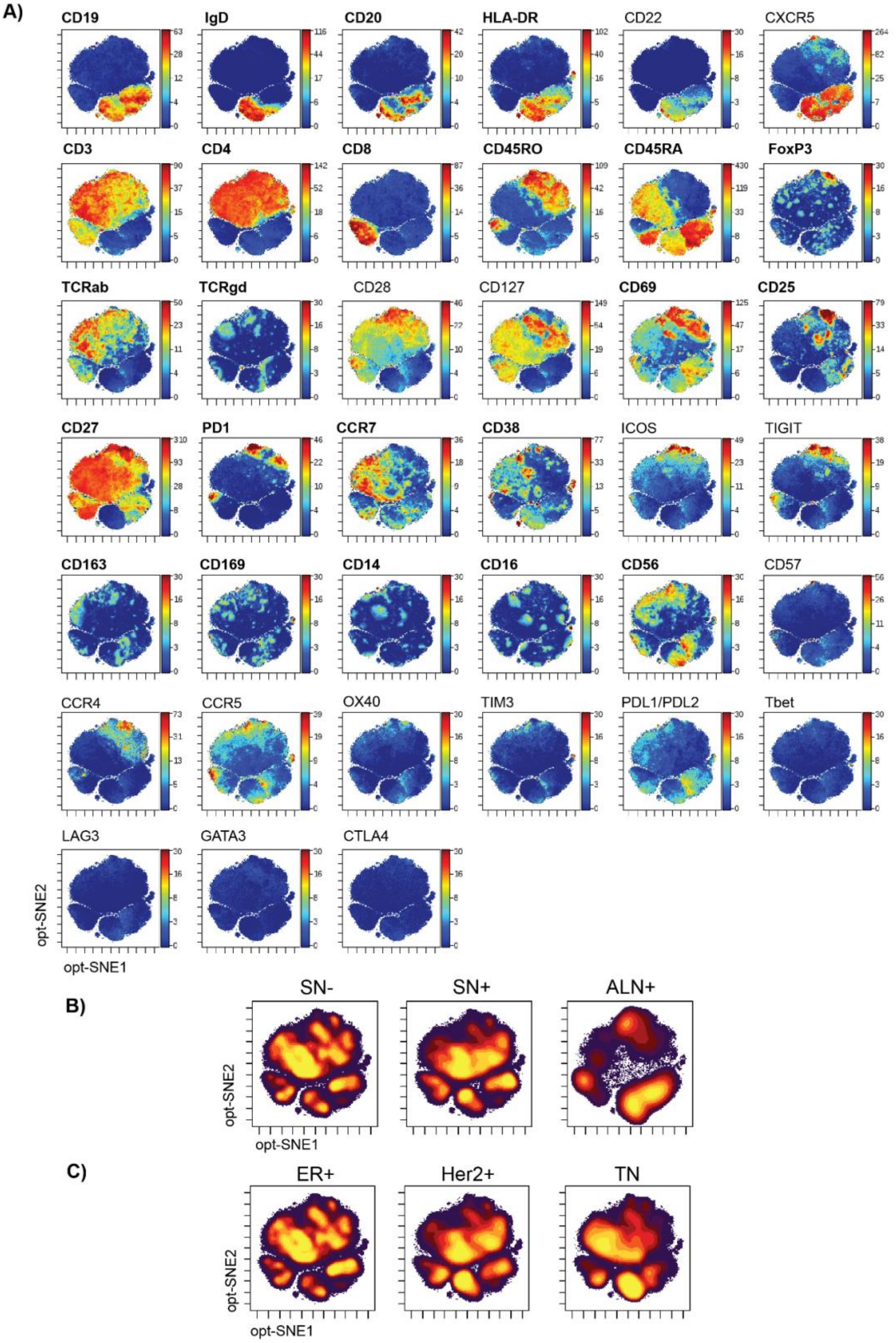


**Supplementary Figure 3:** Expression of immune-cell markers on immune cells from all 458 lymph nodes, visualized by optSNE (markers in bold were included in the optSNE analysis). **A)** All samples concatenated and colored by markers as annotated. **B-C)** Density plot showing the distribution of immune cells across the different lymph node statuses **(B)** and in the three subtypes **(C)**. (SN- = sentinel node without metastasis, SN+ = metastatic sentinel node and ALN+ = metastatic axillary nodes).

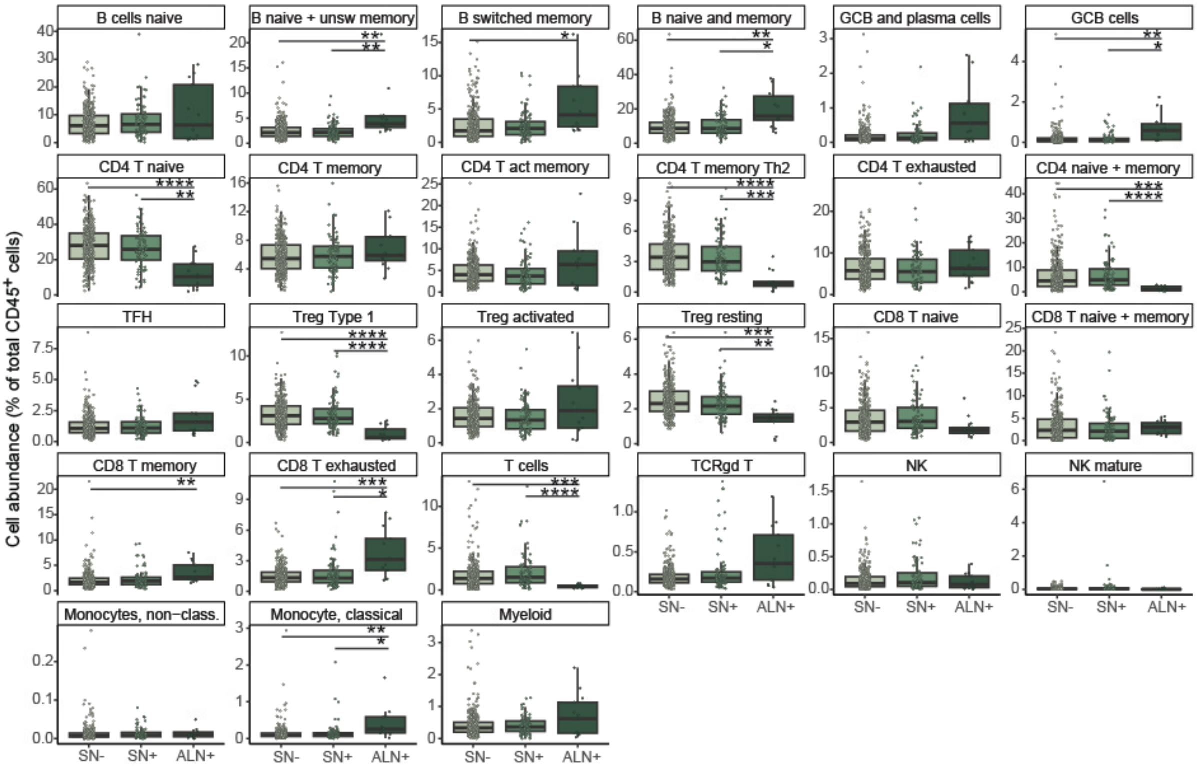


**Supplementary Figure 4**: Differences in cell frequencies across tumor progression within all breast cancer subtypes (SN- = sentinel node without metastasis, SN+ = metastatic sentinel node and ALN+ = metastatic axillary nodes).

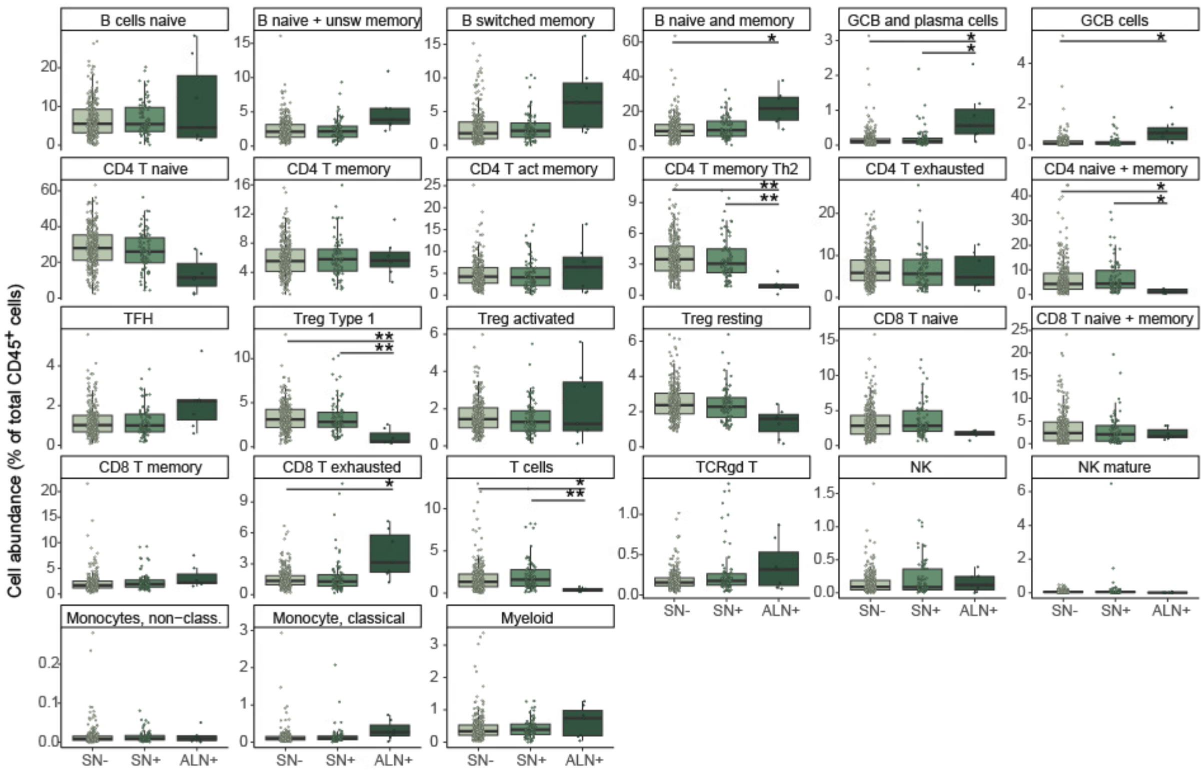


**Supplementary Fig. 5:** Differences in cell frequencies across tumor progression across all samples from ER+ patients (SN- = sentinel node without metastasis, SN+ = metastatic sentinel node and ALN+ = metastatic axillary nodes).

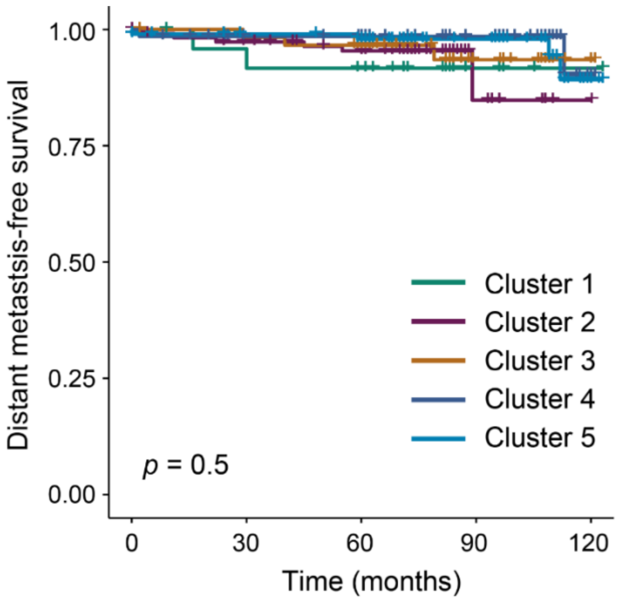


**Supplementary Figure 6**: Kaplan-Meier curves showing disease-free survival (DFS) when stratifying patients according to cluster identity for ER+ samples only. Pairwise log-rank tests with Benjamini-Hochberg post-hoc correction was performed on the survival curves.

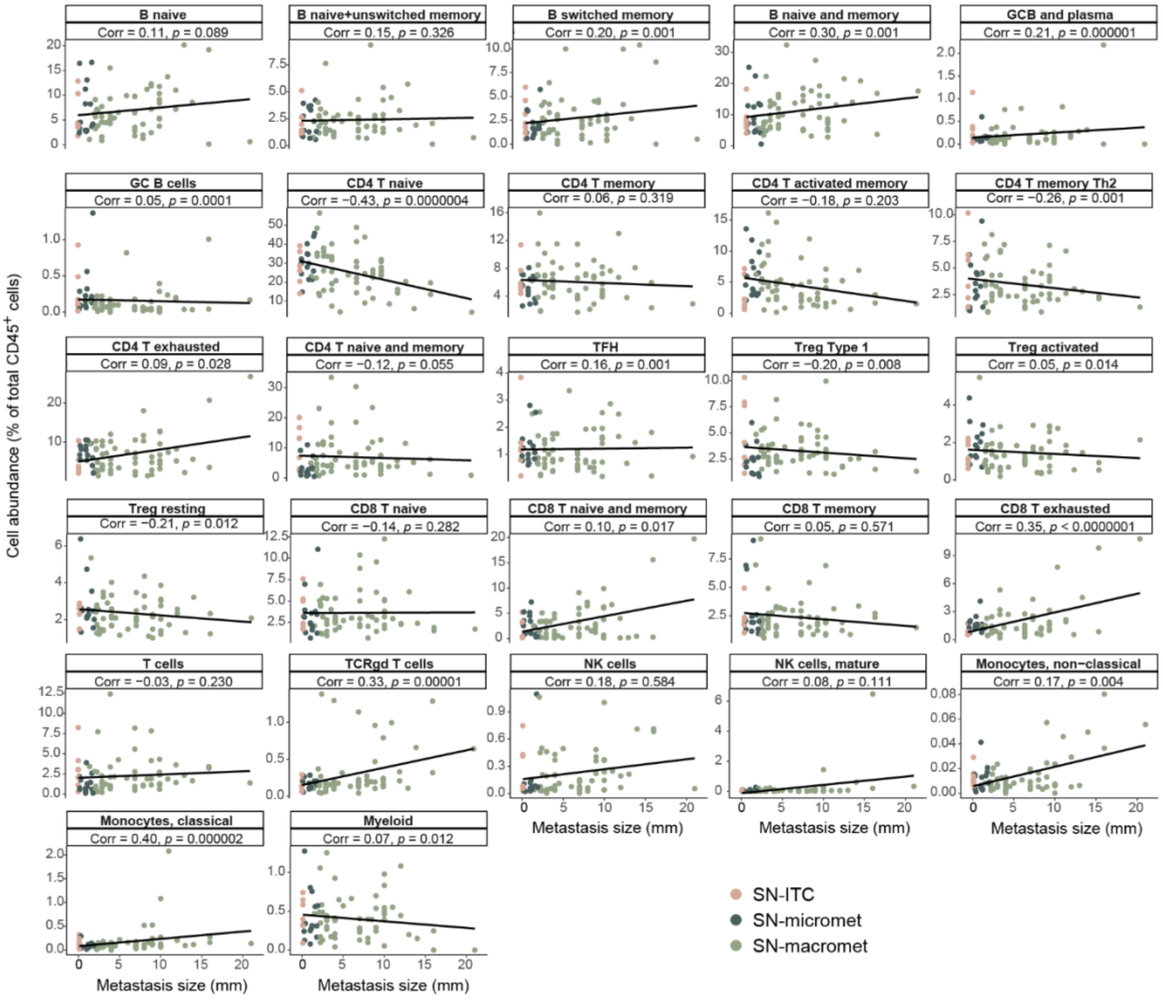


**Supplementary Fig. 7:** Linear regression of metastasis size vs. immune cell populations in SN+ samples. Samples are colored by metastasis size of SN+. (ITC = isolated tumor cells).

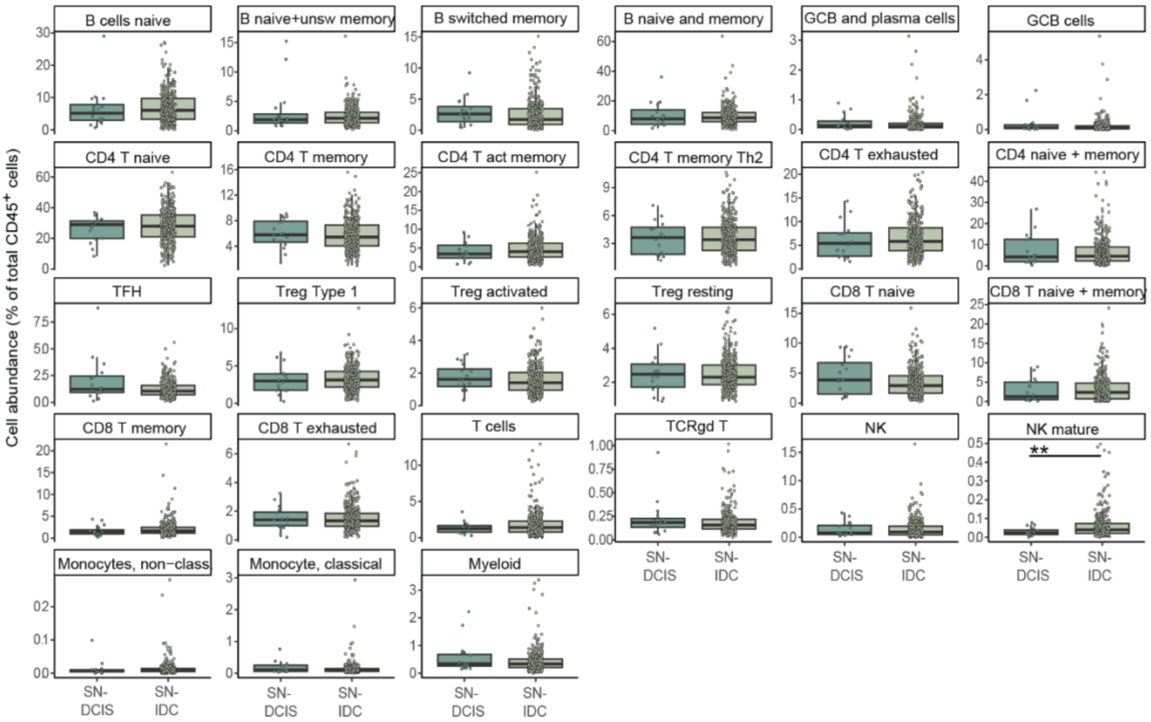


**Supplementary Fig. 8**: **Differences between non-invasive and invasive primary tumors among SN- samples from ER+ patients.** Immune cell frequencies of non-invasive DCIS samples and invasive SN- samples from ER+ patients. Boxes indicate median value and upper and lower quartile, whiskers indicate maximum and minimum data values, and all sample values are shown as dots.

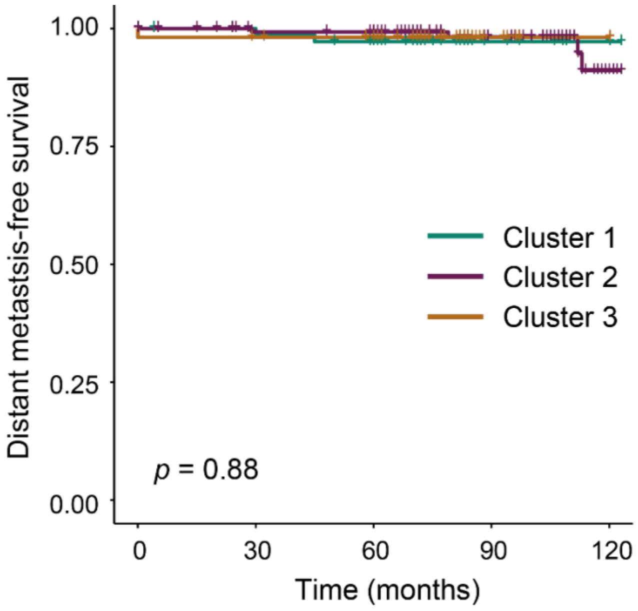


**Supplementary Figure 9**. Kaplan-Meier curves showing disease-free survival (DFS) when stratifying patients according to cluster identity for SN- samples from ER+ patients only (Figure 6). Pairwise log-rank tests with Benjamini-Hochberg post-hoc correction was performed on the survival curves.

**Supplementary Table 1:** Antibodies and open channels in mass cytometry analysis and in FlowSOM analyses.

| Tag | Target | Clone | Vendor | Catalog # | FlowSOM 1 (cleanup) | FlowSOM 2 (cell types) |
|---|---|---|---|---|---|---|
| 89Y | CD45 | HI30 | Standard BioTools | 3089003B | X | X |
| 106Cd | CD22 | HIB22 | Biolegend | 302511 | | X |
| 110Cd | CD3 | UCHT1 | Biolegend | 300443 | | X |
| 111Cd | CD19 | HIB19 | Biolegend | 302247 | | X |
| 112Cd | CD8 | SKI | Biolegend | 344727 | | X |
| 113Cd | Pan Cytokeratin | C11 | Thermo Fisher | Ma1-12594 | X | X |
| 114Cd | CD45RO | UCHL1 | Biolegend | 304239 | | X |
| 116Cd | CD20 | 2H7 | Biolegend | 302343 | | X |
| 138Ba | | | | | X | |
| 139La | | | | | X | |
| 141Pr | EpCAM (CD326) | 9C4 | Standard BioTools | 3141006B | X | X |
| 142Nd | Caspase 3 (Cleaved) | D3E9 | Standard BioTools | 3142004A | X | X |
| 143Nd | CD45RA | HI100 | Standard BioTools | 3143006B | | X |
| 144Nd | CD69 | FN50 | Standard BioTools | 3144018B | | X |
| 145Nd | CD163 | GHI/61 | Standard BioTools | 3145010B | | X |
| 146Nd | CD56 | HCD56 | Biolegend | 318345 | | X |
| 147Sm | TCRab | IP26 | Biolegend | 306702 | | X |
| 148Nd | ICOS (CD278) | C398.4A | Standard BioTools | 3148019B | | X |
| 149Sm | CD25 (IL-2R) | 2A3 | Standard BioTools | 3149010B | | X |
| 150Nd | OX40 (CD134) | ACT35 | Standard BioTools | 3150023B | | X |
| 151Eu | TIM-3 | F38-2E2 | Biolegend | 345019 | | X |
| 152Sm | TCRgd | 11F2 | Standard BioTools | 3152008B | | X |
| 153Eu | CCR4 | L291H4 | Standard BioTools | 3153030A | | X |
| 154Sm | TIGIT | MBSA43 | Standard BioTools | 3154016B | | X |
| 155Gd | CD27 | L128 | Standard BioTools | 3155001B | | X |
| 156Gd | CCR5 | NP-6G4 | Standard BioTools | 3156015A | | X |
| 157Gd | | | | | X | |
| 158Gd | CD169 | 7-239 | Standard BioTools | 3158027B | | X |
| 159Tb | PD-L2 | 24F.10C12 | Biolegend | 329613 | | X |
| 159Tb | CD274 (PD-L1) | 29E.2A3 | Standard BioTools | 3159029B | | X |
| 160Gd | CD28 | CD28.2 | Standard BioTools | 3160003B | | X |
| 161Dy | T-bet | 4B10 | Standard BioTools | 3161014B | | X |
| 162Dy | Foxp3 | PCH101 | Standard BioTools | 3162011A | | X |
| 163Dy | CD57 | HCD57 | Standard BioTools | 3163022B | | X |
| 164Dy | CXCR5 (CD185) | RF8B2 | Standard BioTools | 3164029B | | X |
| 165Ho | LAG3 (CD233) | 11C3C65 | Standard BioTools | 3165037B | | X |
| 166Er | CCR7 | G043H7 | Biolegend | 353237 | | X |
| 167Er | GATA3 | TWAJ | Standard BioTools | 3167007A | | X |
| 168Er | CD127 (IL-7Ra) | A019D5 | Standard BioTools | 3168017B | | X |
| 169Gd | IgD | IA6-2 | Biolegend | 348235 | | X |
| 170Er | CTLA-4 (CD152) | 14D3 | Standard BioTools | 3170005B | | X |
| 171Yb | CD14 | Y1/82A | Standard BioTools | 3171011B | | X |
| 172Yb | CD38 | HIT2 | Standard BioTools | 3172007B | | X |
| 173Yb | HLA-DR | L243 | Standard BioTools | 3173005B | | X |
| 174Yb | CD4 | SK3 | Standard BioTools | 3174004B | | X |
| 175Lu | CD279 (PD-1) | EH12.2H7 | Standard BioTools | 3175008B | | X |
| 176Yb | Histone 3 | D1H2 | Standard BioTools | 3176016A | X | |
| 191/193Ir | Intercalator-Ir | | Standard BioTools | 201192B | X | |
| 194Pt | Cisplatin-194Pt | | Standard BioTools | 201194 | X | |
| 208Pb | | | | | X | |
| 209Bi | CD16 | 3G8 | Standard BioTools | 3209002B | | X |

**Supplementary Table 2**: *p*-values from Mann-Whitney *U* test (after Bonferroni correction, *n* = 27) between all lymph node samples included in Figure 2 and Supplementary Fig. 4.

| **Cell type** | **SN- vs. SN+** | **SN- vs. ALN+** | **SN+ vs. ALN+** |
|---|---|---|---|
| B cells naive | n.s. | n.s. | n.s. |
| B naive and memory | n.s. | 0.003 | 0.021 |
| B naive and unsw mem | n.s. | 0.001 | 0.002 |
| B switched memory | n.s. | 0.015 | n.s. |
| GC B cells | n.s. | 0.002 | 0.012 |
| GCB and plasma | n.s. | n.s. | n.s. |
| CD4 T naive | n.s. | 0.001 | 0.001 |
| CD4 T memory | n.s. | n.s. | n.s. |
| CD4 T activated memory | n.s. | n.s. | n.s. |
| CD4 T naive and mem | n.s. | 0.0001 | 0.0001 |
| CD4 T memory Th2 | n.s. | 0.00001 | 0.0001 |
| CD4 T exhausted | n.s. | n.s. | n.s. |
| TFH | n.s. | n.s. | n.s. |
| Treg Type 1 | n.s. | 0.00001 | 0.00002 |
| Treg activated | n.s. | n.s. | n.s. |
| Treg resting | n.s. | 0.001 | 0.008 |
| CD8 T naive | n.s. | n.s. | n.s. |
| CD8 T naive and memory | n.s. | n.s. | n.s. |
| CD8 T memory | n.s. | 0.010 | n.s. |
| CD8 T exhausted | n.s. | 0.001 | 0.010 |
| T cells | n.s. | 0.0001 | 0.0001 |
| TCRgd T | n.s. | n.s. | n.s. |
| NK cells | n.s. | n.s. | n.s. |
| NK cells, mature | n.s. | n.s. | n.s. |
| Monocytes classical MC2+21) | n.s. | 0.002 | 0.019 |
| Monocytes non-classical | n.s. | n.s. | n.s. |
| Myeloid cells | n.s. | n.s. | n.s. |

**Supplementary Table 3**: *p*-values from Mann-Whitney *U* test (after Bonferroni correction, *n* = 27) between breast cancer subtypes in SN- and SN+ samples, data included in Figure 4.

| | SN- | | | SN+ | | |
|---|---|---|---|---|---|---|
| **Cell type** | ER+ vs. Her2+ | ER+ vs. TN | Her2+ vs. TN | ER+ vs. Her2+ | ER+ vs. TN | Her2+ vs. TN |
| B cells naive | n.s. | n.s. | n.s. | **0.015** | n.s. | n.s. |
| B naive and memory | n.s. | n.s. | n.s. | n.s. | n.s. | n.s. |
| B naive and unsw mem | n.s. | n.s. | n.s. | n.s. | n.s. | n.s. |
| B switched memory | n.s. | n.s. | n.s. | n.s. | n.s. | n.s. |
| GC B cells | n.s. | n.s. | n.s. | n.s. | n.s. | n.s. |
| GCB and plasma | n.s. | n.s. | n.s. | n.s. | n.s. | n.s. |
| CD4 T naive | n.s. | n.s. | n.s. | n.s. | n.s. | n.s. |
| CD4 T memory | n.s. | n.s. | n.s. | n.s. | n.s. | n.s. |
| CD4 T activated memory | n.s. | n.s. | n.s. | n.s. | n.s. | n.s. |
| CD4 T naive and mem | n.s. | n.s. | n.s. | n.s. | n.s. | n.s. |
| CD4 T memory Th2 | n.s. | n.s. | n.s. | n.s. | n.s. | n.s. |
| CD4 T exhausted | n.s. | n.s. | n.s. | n.s. | n.s. | n.s. |
| TFH | n.s. | n.s. | n.s. | n.s. | n.s. | n.s. |
| Treg Type 1 | n.s. | n.s. | n.s. | n.s. | n.s. | n.s. |
| Treg activated | n.s. | n.s. | n.s. | n.s. | n.s. | n.s. |
| Treg resting | n.s. | n.s. | n.s. | n.s. | n.s. | n.s. |
| CD8 T naive | n.s. | n.s. | n.s. | n.s. | n.s. | n.s. |
| CD8 T naive and memory | n.s. | n.s. | n.s. | n.s. | n.s. | n.s. |
| CD8 T memory | n.s. | n.s. | n.s. | n.s. | n.s. | n.s. |
| CD8 T exhausted | n.s. | n.s. | n.s. | n.s. | n.s. | n.s. |
| T cells | n.s. | n.s. | n.s. | n.s. | n.s. | n.s. |
| TCRgd T | n.s. | n.s. | n.s. | n.s. | n.s. | n.s. |
| NK cells | n.s. | n.s. | n.s. | n.s. | n.s. | n.s. |
| NK mature | n.s. | n.s. | n.s. | n.s. | n.s. | n.s. |
| Monocytes classical | n.s. | n.s. | n.s. | n.s. | n.s. | n.s. |
| Monocytes non-classical | n.s. | n.s. | n.s. | n.s. | n.s. | n.s. |
| Myeloid cells | n.s. | n.s. | n.s. | n.s. | n.s. | n.s. |

**Supplementary Table 3**: *p*-values from Mann-Whitney *U* test (after Bonferroni correction, *n* = 27) between breast cancer subtypes in lymph node samples from ER+ patients included in Figure 5 and Supplementary Fig. 5.

| **Cell type** | **SN- vs. SN+** | **SN- vs. ALN+** | **SN+ vs. ALN+** |
|---|---|---|---|
| B cells naive | n.s. | n.s. | n.s. |
| B naive and memory | n.s. | 0.014 | n.s. |
| B naive and unsw mem | n.s. | n.s. | n.s. |
| B switched memory | n.s. | n.s. | n.s. |
| GC B cells | n.s. | 0.034 | n.s. |
| GCB and plasma | n.s. | 0.015 | 0.048 |
| CD4 T naive | n.s. | n.s. | n.s. |
| CD4 T memory | n.s. | n.s. | n.s. |
| CD4 T activated memory | n.s. | n.s. | n.s. |
| CD4 T naive and mem | n.s. | 0.025 | 0.019 |
| CD4 T memory Th2 | n.s. | 0.001 | 0.003 |
| CD4 T exhausted | n.s. | n.s. | n.s. |
| TFH | n.s. | n.s. | n.s. |
| Treg Type 1 | n.s. | 0.006 | 0.008 |
| Treg activated | n.s. | n.s. | n.s. |
| Treg resting | n.s. | n.s. | n.s. |
| CD8 T naive | n.s. | n.s. | n.s. |
| CD8 T naive and memory | n.s. | n.s. | n.s. |
| CD8 T memory | n.s. | n.s. | n.s. |
| CD8 T exhausted | n.s. | 0.030 | n.s. |
| T cells | n.s. | 0.016 | 0.009 |
| TCRgd T | n.s. | n.s. | n.s. |
| NK cells | n.s. | n.s. | n.s. |
| NK mature | n.s. | n.s. | n.s. |
| Monocytes classical | n.s. | n.s. | n.s. |
| Monocytes non-classical | n.s. | n.s. | n.s. |
| Myeloid cells | n.s. | n.s. | n.s. |

**Supplementary Table 4:** *p*-values from Mann-Whitney *U* test (after Bonferroni correction, *n* = 27) between different clinical parameters in SN- samples from ER+ patients.

| | **Invasiveness** | **Tumorsize** | **Ki67** | **Age** | **Tumor grade** |
|---|---|---|---|---|---|
| **Cell type** | DCIS vs. IDC | Small vs. Large (median = 50 mm) | High vs. Low (median = 26%) | Above vs. below 50 years | 1 vs. 2, 1 vs. 3, 2 vs. 3 |
| B naive | n.s. | n.s. | n.s. | n.s. | n.s. |
| B naive and memory | n.s. | n.s. | n.s. | n.s. | n.s. |
| B naive and unsw mem | n.s. | n.s. | n.s. | n.s. | n.s. |
| B switched memory | n.s. | n.s. | n.s. | n.s. | n.s. |
| GC B cells | n.s. | n.s. | n.s. | n.s. | n.s. |
| GCB and plasma | n.s. | n.s. | n.s. | n.s. | n.s. |
| CD4 T naive | n.s. | n.s. | n.s. | n.s. | n.s. |
| CD4 memory | n.s. | n.s. | n.s. | n.s. | n.s. |
| CD4 activated memory | n.s. | n.s. | n.s. | n.s. | n.s. |
| CD4 naive and mem | n.s. | n.s. | n.s. | n.s. | n.s. |
| CD4 memory Th2 | n.s. | n.s. | n.s. | n.s. | n.s. |
| CD4 T exhausted | n.s. | n.s. | n.s. | n.s. | n.s. |
| TFH | n.s. | n.s. | n.s. | n.s. | n.s. |
| Treg Type 1 | n.s. | n.s. | n.s. | n.s. | n.s. |
| Treg activated | n.s. | n.s. | n.s. | 0.022 | n.s. |
| Treg resting | n.s. | n.s. | n.s. | n.s. | n.s. |
| CD8 naive | n.s. | n.s. | n.s. | 0.000002 | n.s. |
| CD8 naive and mem | n.s. | n.s. | n.s. | n.s. | n.s. |
| CD8 memory | n.s. | n.s. | n.s. | n.s. | n.s. |
| CD8 exhausted | n.s. | n.s. | n.s. | n.s. | n.s. |
| T cells | n.s. | n.s. | n.s. | n.s. | n.s. |
| TCRgd T | n.s. | n.s. | n.s. | n.s. | n.s. |
| NK cells | n.s. | n.s. | n.s. | n.s. | n.s. |
| NK cells, mature | n.s. | n.s. | n.s. | n.s. | n.s. |
| Monocytes classical | n.s. | n.s. | n.s. | n.s. | n.s. |
| Monocytes non-classical | n.s. | n.s. | n.s. | n.s. | n.s. |
| Myeloid | n.s. | n.s. | n.s. | n.s. | n.s. |

**Supplementary Table 5:** Number of samples with tumor cells detected by mass cytometry (CyTOF) analysis in SN-, SN+ and ALN+ lymph nodes. Samples were defined as tumor-positive when tumor cells were >0.02% of total cells.

| | **Tumor-positive** | **Tumor-negative** |
|---|---|---|
| **SN-** | 73 (20.4%) | 284 (79.6%) |
| **SN+** | 42 (47.7%) | 46 (52.3%) |
| **ALN+** | 9 (69.2%) | 4 (30.8%) |
